# Adaptative Machine Translation between paired Single-Cell Multi-Omics Data

**DOI:** 10.1101/2021.01.27.428400

**Authors:** Xabier Martinez-de-Morentin, Sumeer A. Khan, Robert Lehmann, Sisi Qu, Alberto Maillo, Narsis A. Kiani, Felipe Prosper, Jesper Tegner, David Gomez-Cabrero

## Abstract

**Background:** Single-cell multi-omics technologies allow the profiling of different data modalities from the same cell. However, while isolated modalities only capture one view of the total information of a biological cell, an integrative analysis capturing the different modalities is challenging. In response, bioinformatics and machine learning methodologies have been developed for multi-omics single-cell analysis. Nevertheless, it is unclear if current tools can address the dual aspect of modality integration and prediction across modalities without requiring extensive parameter finetuning.

**Results:** We designed LIBRA, a Neural Network based framework, to learn a translation between paired multi-omics profiles such that a shared latent space is constructed. LIBRA is a state-of-the-art tool when evaluating the ability to increase cell-type (clustering) resolution in the latent space. When assessing the predictive power across data modalities, LIBRA outperforms existing tools. Finally, considering the importance of hyperparameters, we implemented an *adaptative-tuning* strategy, labelled aLIBRA, in the LIBRA package. As expected, adaptive parameter optimization significantly boosts the performance of learning predictive models from paired datasets. Additionally, aLIBRA provides parameter combinations balancing the integrative and predictive tasks.

**Conclusions:** LIBRA is a versatile tool, uniquely targeting both integration and prediction tasks of Single-cell multi-omics data. LIBRA is a data-driven robust platform that includes an adaptive learning scheme. Furthermore, LIBRA is freely available as R and Python libraries (https://github.com/TranslationalBioinformaticsUnit/LIBRA).

## Introduction

Single Cell Genomics technologies set the stage for unraveling the intrinsic complex organization at single-cell resolution by simultaneously profiling several layers of transcriptional regulation^1,2,3^. Recent multi-omics single-cell technologies enable profiling of joint “chromatin accessibility & mRNA profiles” (e.g., SNARE-seq^4^, sci-CAR^5^, SHARE-seq^6^, Paired-seq^7^, 10X Genomics^8^), “mRNA profiles & protein antibody-derived tags” (CITE-seq^9^), and even more than omics such as “chromatin accessibility, DNA methylation, and transcriptome profiling” in the scNMT-seq^10^ protocol. As a result of such novel technologies, it became necessary to develop methods to integrate multiomic profiles at single-cell level^11^ (see Fig.1A). The rationale is that current state-of-the-art bulk methodologies^12,13,14,15^ and frameworks^16^ could not analyze single-cell data optimally^17,18,19^. Initially, methodologies such as Seurat3^20^ allowed integrative analysis; however, Seurat3^20^ does not use the information derived from the *paired* nature of the data (*profiles obtained from the same cell*). More recently, methodologies making use of the paired information were developed^21^. For example, machine learning tools such as MOFA+^22^ and Seurat4^23^ allow the identification of an integrated space that can be used for improved cell clustering. However, such tools have two potentially limiting factors: scalability and robustness. Furthermore, they do not allow predicting profiles between omics. Deep Learning-based methodologies were developed to overcome such limitations. The first one was BABEL^24^ which was aimed to generate predictive models “*translating*” between data types. Others followed this idea, such as KPNNs^25^, GAT^26,^ or scNym^27^. Recently, a Multi-modal Single-Cell Data Integration Competition^28^ (denoted as the NeurIPS challenge) was launched, where several Deep Learning (DL) techniques were developed and evaluated. The NeurIPS challenge addressed several tasks such as (a) predicting one modality from another (*prediction*), (b) matching cells between modalities, and (c) jointly learning representations of cellular identity (*representation*). In general, neural networks were the most popular and provided – in most cases - the best results. However, while the best methodologies used architectures of limited complexity, it became apparent during the competition that the methodologies required extensive fine-tuning of hyperparameters regardless of the specific architecture. Furthermore, by observing that no method could win in more than one task, it was concluded that “*no free lunch*” ^29^ (*no method works best for all*) also applies to the multi-omic analysis. Hence, in order to avoid multiple analyses with different tools, methods competitive in several of the tasks are required.

**Figure 1.**
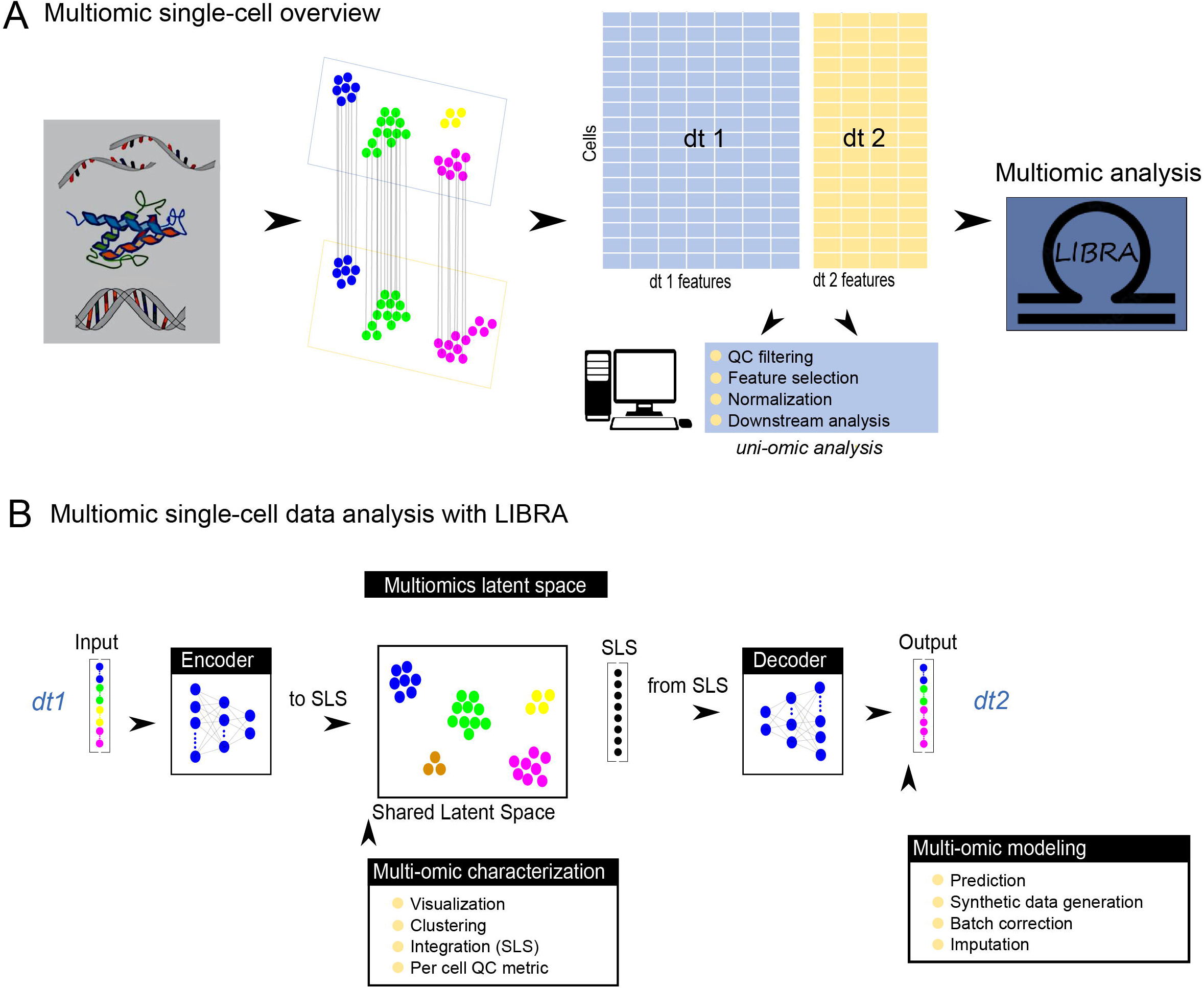
LIBRA overview. **A**. The upper panel provides a visual representation of the task to be solved. From left to right, the initial raw data sets (omics) are represented. Capturing the single cell information from exact same cells from each omic leads to refining the unique provided information from a unique omic. This information needs to be integrated correctly without loss of omics uniqueness information. This requires a preprocessing and integration method. LIBRA is proposed for this end. **B**. The main idea of LIBRA is represented in the lower panel. Through an encoding-decoding process, the information of the cells from respective omics will be extracted and their distance will be minimized while preserving the biological information to generate a latent space that contains the information from both. Additional steps will be taken such as visualization, clustering or prediction. Several measurements are carried out for quality and performance.

Therefore, we propose LIBRA (Fig.1B), an encoder-decoder architecture using AutoEncoders (AE) that can competitively perform the two of the three tasks addressed in the NeurIPS challenge (*prediction* and *representation*). LIBRA is inspired by the ideas from Neural Machine Translation^30^. Similar to BABEL^24^, LIBRA integrates single-cell multi-omics data by leveraging *paired* single-cell omics data. Initially, we fine-tune the first version of LIBRA using a step-wise optimization strategy considering AE-associated quality measures. To this end, we developed a novel metric, Preserved Pairwise Jaccard Index (PPJI), which investigates if the integrated space allows for finer granularity in cell sub-type detection. Interestingly, we observe that PPJI is a valuable metric to quantify the added value of a multi-omic joint *representation*. Secondly, we compare LIBRA with the current state-of-the-art tools for several datasets, data modality combinations, and the different tasks. LIBRA compares competitively in all cases. It is relevant to notice that LIBRA was among the top 10 in two of the NeurIPS challenges (jointly learning and predicting modality) according to the NeurIPS challenge’s metrics. Finally, to address the *no-free-lunch* observation, we combine LIBRA with a self-tuning paradigm^31^ that allows LIBRA parameter optimization on-the-run. Such self-tuning not only improves the results significantly and outperforms available methodologies but also provides instantiations of LIBRA that are competitive at both integration and prediction. LIBRA is freely available in both R and Python (including tutorials) for multi-modal single-cell analysis.

## Materials and Methods

### Preprocessing of sc-RNA-seq data

Following Seurat guidelines, a four-step cell and feature quality filtering were applied. First based on lower (0.1) - upper (0.9) bound quantile for “number of features” and “counts per cell”, followed by a minimum feature per cell filtering allowing cells present in at least 201 genes and a feature removing strategy based on “minimum features per cell”, retaining only genes present in at least 4 cells. Finalizing a cell cut-off base on maximum mitochondrial percentage was applied, allowing cells containing less than 5% mitochondrial gene content. Feature selection criteria have been used for downstream analysis based on most variable genes, making use of 2.000 top genes. Normalization of the gene expression measurements for each cell was done by the total expression and multiplying this by a scale factor (10,000) and log-transforming it. Feature subspaces were obtained from most variable genes using principal component analysis (PCA) using 15 components. Clustering was computed using the Louvain algorithm over principal components subspace. Bootstrap subsampling *snakemake workflow* was used to identify the optimal number of nearest neighborhoods and the resulting resolution. The range of the values for these parameters were 8-16 and 0.6-1.4, respectively. We used a subsampling rate of 0.8 for 20 subsamples, which generated a total of 500 samples for analysis. Clustering was repeated 1.000 times with final settings for excluding additional spurious clusters due to starting seed initialization. Specific actions employed over any datasets are available in Supplementary Materials and Methods section.

As a result, a robust latent space and clustering results are obtained for using as reference sc-RNA-seq to compute performance metrics later. A normalized scRNA-seq matrix will serve as input to the LIBRA model.

### Preprocessing of sc-ATAC-seq data

Following the scRNA-seq schema, sc-ATAC-seq data was also preprocessed similarly except in some cases described below. In this case, a combined Seurat and Signac guideline was used.

Due to increased sparsity of sc-ATAC-seq the minimum features per cell filtering were changed, selecting peaks in at least three cells. Unlike in sc-RNA-seq, data was normalized using the frequency-inverse document frequency (TF-IDF) method. Reducing the features to a list of most variable peaks is an option that may reduce the performance. Instead, we used the full feature space in this case. Reduced feature subspaces were computed over all peaks features space using singular value decomposition (SVD), providing latent semantic indexing (LSI) as latent space with 50 components. Values and functions employed are available in the Supplementary Materials and Methods section.

Aiming to perform integration reference by using Seurat protocols, sc-ATAC-seq preprocessed data was used for generating sc-RNA-simulated data; to this end, the Signac *activity* estimation approach was used. An upstream of 2.000 base pairs was used for “peak to gene relation” estimation, and GRCH38-mm10 reference genomes were used for corresponding datasets specie for human and mouse, respectively. See Supplementary Materials and Methods for Seurat integration information.

As for scRNA-seq, the reduced latent space and clustering results obtained are used as scATAC-seq reference for later performance metrics computation. A normalized scATAC-seq matrix will serve as input to the LIBRA model.

### Preprocessing of CITE-seq data

The initial pipeline for the analysis of CITE-seq raw data was similar to previous data modalities. Differences are detailed below.

The entire protein space was used instead of selecting for most variable proteins sub-space. In addition, normalization of the protein expression measurements for each cell was done using centred log-ratio transformation (CLR). Values and functions employed are available in the Supplementary Materials and Methods section. The reduced latent space and clustering results obtained are used as antibody-derived tags (sc-ADT) reference for later performance metrics computation. A normalized scADT matrix will serve as input to LIBRA model.

## Algorithm & Implementation

### LIBRA framework

LIBRA “*translates”*^15^ between omics. Implemented using Autoencoders, LIBRA encodes one omics and decodes the other omic *to and from* a reduced space. Here the decoder minimizes the distance to a second and paired data type (*joined translation and projection*). Briefly, LIBRA consists of two neural networks (NN) (Fig.2A); the first NN (NN1) is designed similarly to an Autoencoder, but the difference is that input (dt1) and output (dt2) data correspond to two modalities of a paired multi-modal dataset (FigS1A).

**Figure 2.**
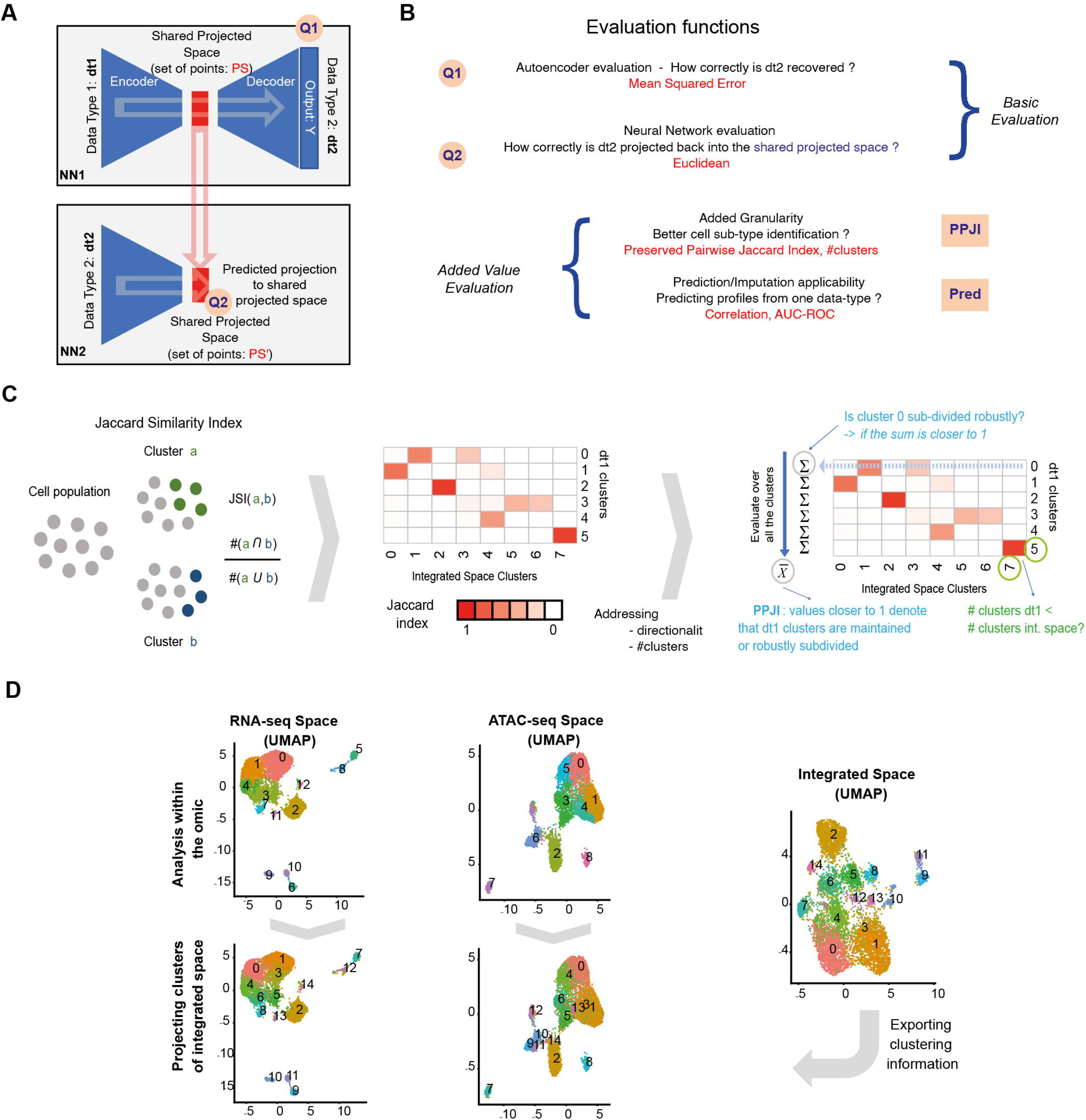
LIBRA design and challenge resolution. **A**. Visual description of the LIBRA framework. LIBRA consists of two neural networks (NN); the first NN (NN1) is designed similarly to an Autoencoder, but the difference is that input (dt1) and output (dt2) data correspond to two modalities of a paired multi-modal dataset. The idea is to learn a shared latent space for two data-types, as shown in panel (a). The second NN maps *dt2 to the shared latent space* to ensure that the projected space correctly embeds the dt2 information. See Fig.S2(a,b) for implementation details after fine-tuning. **B**. Summary overview of the evaluation functions used in the analysis and optimization of LIBRA. See Material and Methods for complete details. **C**. Visual description of the PPJI. Left panel: visual description of the Jaccard Similarity Index. Middle panel: visual description of the Pairwise Jaccard Similarity Index. Right panel: visual description of the Preserved Pairwise Jaccard Index (PPJI); as shown in the figure PPJI investigates if the clusters derived from a single-omic data-analysis (dt1) are properly separated robustly into the same or larger number of clusters. To this end, for each cluster i derived from dt1, the sum of JSI(*i,x*) for all clusters *x* in the integrated space is computed. And then, the average for all clusters from dt1 is computed as the final summary. A value of 1 denotes that dt1 clusters are perfectly identified in the integrated space; however, an added value requires large values of PPJI but also a larger number of clusters identified in the integrated space. An extended description of PPJI computation is provided in the methods section. **D**. Example of the integrative challenge using dataset DS1. The integrated space was identified using LIBRA. Example of clustering resolution in the integrated space. Two left upper panels denote the UMAP projection and clustering for RNA and ATAC, respectively. The right panel shows the UMAP projection and clustering of cells in the integrated space (e.g., the LIBRA optimized model). Finally, the two left bottom panels project the clustering information derived from the integrated space in the UMAP projections for RNA and ATAC, respectively.

Considering only one hidden layer, the encoder part of NN1 will aim to encode the input omic expression matrix detonated as x ∈ ℝ^*d*^ to the latent variables *h* following this formula:

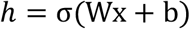

Where σ is the element-wise activation function, *W* is the weight matrix, and *b* is the bias vector. In LIBRA implementation activation function used is leaky-relu ^32^, this decision was taken due to a high rate of dead nodes generation using standard relu, producing lower performance and uncertainty on outcomes delivery. The activation function is then, instead of being 0 when *z* < 0, a small non-zero, constant gradient *α* where function is as follows 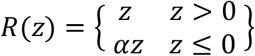 and its derivative as 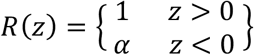. This selection excludes LIBRA training to introduce death nodes due to the sparsity nature of the single-cell data. Weights and biases are initialized using normal distributed criteria *Xavier uniform initializer* and *zeros*. The encoding part of the autoencoder follows this formula:

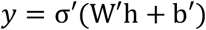

Where output omic expression matrix detonated as y ∈ ℝ^*d*^ it’s used to force the loss function to be minimized based on a second omic instead of the original input. This process will be repeated during training using backpropagation for weight updating. The loss function employed is the mean squared error (MSE). An early stopping rule was added to save time when the evaluation function cannot retrieve better scores for MSE with a fixed patience value. In addition, a learning rate plateau callback was added to benefit from reducing the learning rate when no improvements are obtained on loss function during a fixed patience value. See Supplementary Materials and Methods for values and hyperparameters employed.

Thanks to this processing, a shared latent space (SLS) for two data types can be learned effectively (Fig.2A). While NN1 identifies the SLS, we considered it necessary to implement a second NN(NN2) that maps *dt2* to the generated SLS to ensure and quantify that the projected space correctly embeds the dt2 cells information with a high quality (Fig.2D). This NN2 (Fig.S1B) will use the same encoding strategy but for *dt2* as input but will contain generated SLS as output following this formula:

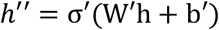

Where *h*^″^ is the encoded SLS generated by NN1.

### Evaluation functions

To evaluate the integration performance of LIBRA, we designed several quality metrics (Fig.2B,C). The first set of metrics, Q1 and Q2 (Fig.2B, upper part), are associated with the neural network training; the mean square error (MSE^33^) and the Euclidean distance are used to evaluate the training of NN1 and NN2, respectively. The second set of metrics (Fig.2B, lower part) was implemented to evaluate the applications of LIBRA: added value of integration and predictive power between omic profiles. The following sub-sections describe each of the evaluation functions; for additional technical details, see Supplementary Information.

#### Q1 has been computed as the MSE formula for NN1

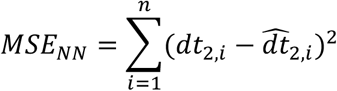

Where 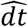 denotes the estimated value, and *dt* denotes the original value. *n* is the total number of cells for the given pair of single-cell data modalities.

#### Q2 as the Euclidean distances between generated SLS

As NN2 is trained using MSE as a loss function, the Q2 Euclidean distance between SLS generated in NN1 and output from NN2 predicted values could easily be computed as the root square of MSE obtained in NN2 when computed for all cells as:

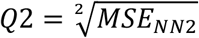

In the case of NN2 and based on the observed bimodal distribution (Fig.S1C), we evaluated separately the cells used in the training or the validation in NN1.

#### Preserved Pairwise Jaccard Index

The Preserved Pairwise Jaccard Index (PPJI) is designed to quantify the added value of the paired integrative analysis to identify cell subtypes (better granularity) by providing a summary value over the Pairwise Jaccard Index (PJI) matrix (Fig.2C). Briefly, PPJI provides a number between 0 and 1 quantifying: “*does the integrated space provide a finer cell-type definition than the cell-type definitions generated from a single-omics (e*.*g*., *dt1)*?”. For a given cluster in dt1, the sum over the PJI matrix associated will be “1” if the cluster is maintained or perfectly separated into sub-clusters (Fig.1C). Thus, PPJI computes the average of the sums as a summary. PPJI computation will be as follows (See Supplementary Materials and Methods for detailed explanation):

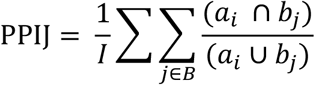

Where for every pair of clusters i ∈ A and *j* ∈ B (a_i_ and b_*j*_ denote the set of cells in cluster *i* and *j* respectively) when aiming to investigate how A clustering projects on B clustering of reference (see Fig.2D). Consequently, values closer to one denote that clusters in dt1 are either preserved or further split into sub-clusters. Importantly, PPJI should be combined to compare the number of clusters in the dt1 and the integrated space.

#### Synergy model performance ranking

For a summary score for the integration performance, a weighted average was computed over the three metrics (See Supplementary Materials and Methods, See Table S2); where each metrics is scaled and weighted equally. To that end, every time a set of combinations are compared, the results of training the AE 10 times for each combination are grouped. Then for Q1 (Q2), the maximum (*maxQ1*) and minimum (*minQ1*) are computed. For each running value of *I*, a *snormQ1* (*snormQ2*) is computed as follows:

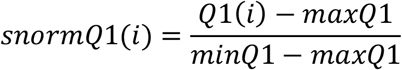

Higher values are associated with a lower error. For PPJI, considering the aim is to maximize the value, the values are computed as follows:

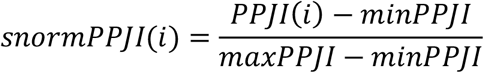

The weighted combination (*score*) for network training *i* is computed as follows:

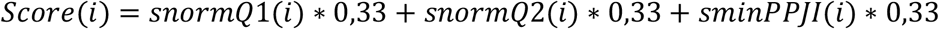

Larger *score* values denote a better performance, generally associated with lower Q1 and Q2, and higher PPJI values.

#### Prediction specific evaluation metrics

To evaluate the predictive power of LIBRA, the *pred* metric is used. The Pearson correlation and AUC-ROC curves were used for scRNA-seq and scATAC-seq, respectively. For ROC computation scATAC-seq predicted matrix was first binarized employing a 0.25 value as cut-off (based on the data distribution): values larger than 0.25 are considered as 1, values below or equal to 0.25 are considered as 0. For details of the implementation of this metric, see Supplementary Materials and Methods.

#### CITE-seq specific integration measurement

The last metric implemented is a CITE-seq^9^ specific for integration performance measurement. For this metric, a set of 25 reference expression proteins are used to measure how different are the Spearman and Pearson correlation scores for the k-nearest neighbouring cells (k=20) on the reference protein dataset entire feature space for each of these proteins to the expression of the k-nearest neighbouring cells obtained in the SLS for the different methods employed (LIBRA, Seurat4^23^, MOFA+^22^, totalVI^34,^ and BABEL^24^) for each the 25 reference proteins.

### Adaptative fine-tuning, aLIBRA

The first version of the LIBRA framework was instantiated using a step-wise optimization procedure over a single dataset; as a result, a set of parameters were selected, and such a framework was applied to all datasets. Such step-wise optimization is detailed in the Results section. However, such parameter combinations may work sub-optimally for different datasets or combinations of data modalities. Based on that assumption, we combine LIBRA with an automatic grid-based fine-tuning strategy to identify the optimal set of parameters for any dataset; we denote the implementation as **adaptative LIBRA** (aLIBRA).

In aLIBRA, the optimal combination of the *number of layers, number of nodes, alpha, dropout*, and *mid-layers size* for NN1 and NN2 is identified. A non-linear decrease has been used for the hidden layer size rule generation following *Layer size* _*N =*_ *input layer size /2 * N* for both encoding and the reverse augmentation on the decoding part of the autoencoder; N denotes the position a layer has in a neural network. Such consideration prevents LIBRA from generating smaller layers than the size of the “middle layer” for very large NN (that may be required in very large datasets).

The fine-tuning is run twice for each of the tasks: integration and prediction. For integration, aLIBRA considered the following values: number of layers [1,2,3,4,5,6], number of nodes [256,512,1024,2048], alpha [0.1,0.3,0.5], dropout [0.1,0.2,0.3,0.4] and mid layers size [10,50,70]. For prediction, aLIBRA considered the following values: number of layers [1,2], number of nodes [128,256,512], alpha [0.05,0.1,0.3], dropout [0.1,0.2], batch size [32,64,128] and mid layers size [10,30,50,70]. Those options are customizable in the Python implementation.

The aLIBRA fine-tuning has been implemented with a parallelization strategy to decrease the computation time requirements. For further details, see Supplementary Materials and Methods.

## Results

### LIBRA step-wise optimization

To identify default-tuned LIBRA’s hyperparameters, we combined three quality measurements (Q1, Q2, PPJI) and the SNARE-seq^4^ adult brain mouse dataset (DS1). Iteratively, we considered the optimization of the following parameters: (i) Autoencoder-type configuration=**AE-based framework**, (ii) number of dimensions of the projected space=**10**, (iii) peak derived information for ATAC-seq, (iv) the ordering (**dt1=ATAC-seq and dt2=RNA-seq**), (v) to consider **most variable features** only, and (vi) the number of hidden layers=**2**. Table S1 includes the values for each evaluation metric. In all cases, a weighted *score* combining Q1, Q2, and PPJI was computed to determine the overall performance. Table S2 shows the final weighted *score* computed for each running in each combination. The best configuration was chosen due to overrepresentation within the 10 highest values on each parameter selection step. See additional details in Supplementary Materials and Methods).

### Comparing LIBRA with existing tools

Next, we compared step-wise fine-tuned LIBRA using DS1 against existing tools Seurat3^20^, Seurat4^23^, MOFA+^22^, totalVI^34,^ and BABEL^24^. For that comparison we used the PPJI measure (Fig.2C,D) which quantifies the added value of multi-omic integration when identifying cell sub-types (Fig.3A). We observed that only Seurat4^23^ minimally outperforms default-tuned LIBRA. Yet, inspecting the clusters reveals an overwhelming similarity, as shown in Fig.S1D,E, suggesting that the minor quantitative difference is not critical from a biological standpoint. Interestingly, LIBRA outperforms the other deep learning frameworks including a concatenation of both data-type matrices in an Autoencoder to identify the shared latent space (*unpaired AE*). We investigated the cluster-specific markers from Seurat4^23^ and LIBRA to interrogate biological relevance. First, when taking Seurat4^23^ as a reference, the top markers identified at Seurat4^23^ are also recognized by LIBRA (Fig.S2A,B). LIBRA also identifies other markers (Fig.S2A,B). We conclude that both methodologies can recover a similar level of resolution for clusters, cell subtypes, and their associated biomarkers.

**Figure 3.**
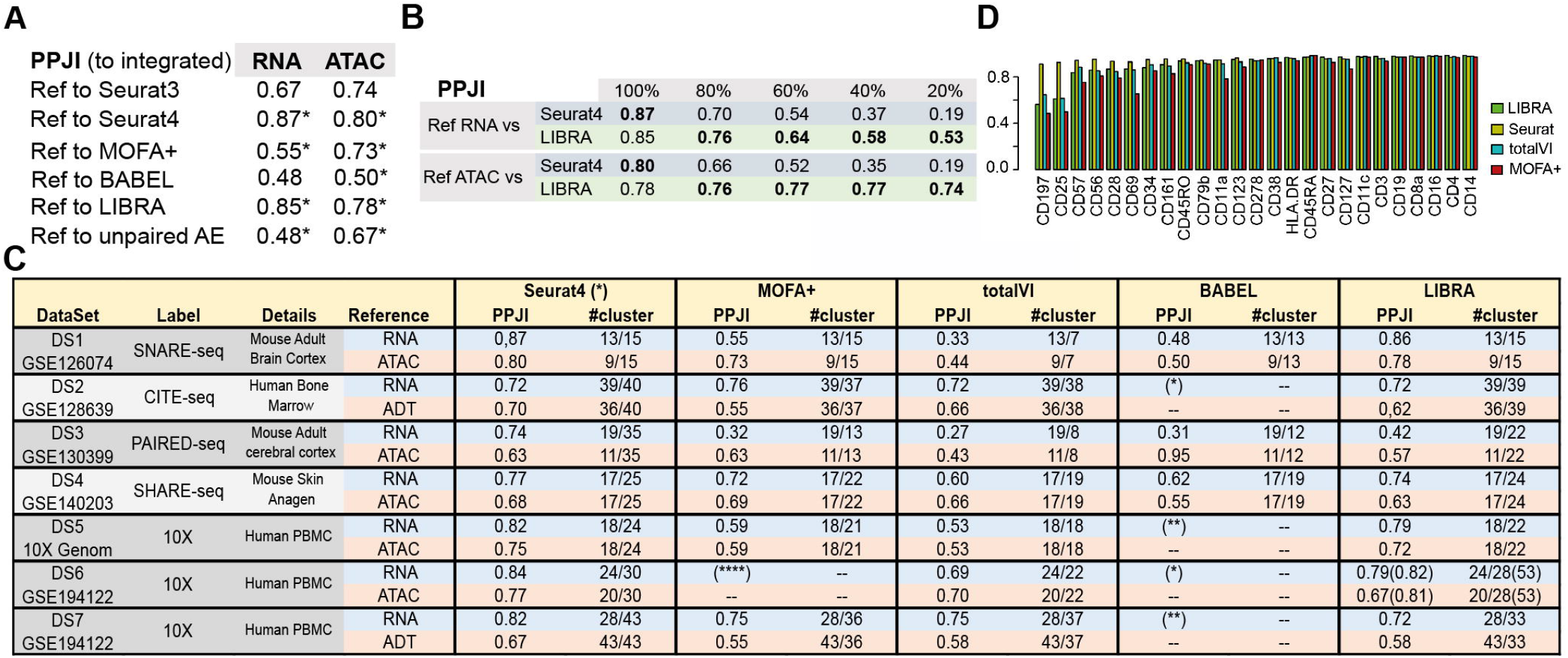
LIBRA step-wise tuning and comparison with existing methodologies. **A**. PPJI evaluation in DS1 for each of the methodologies considered in the analysis. * Indicates that the number of clusters in the integrated space is larger. **B**. PPJI values derived from LIBRA and Seurat4, both using paired information when the total number of cells used to create the model is a percentage from the original total number (6735 cells). **C**. PPJI-based comparison between the different methodologies in several datasets. (*) Not conducted in BABEL. (**) DS5 was analyzed using all genes instead of MVG. BABEL framework exceeded the limiting time of 1 week for running on a GPU infrastructure. (***) DS6 was analyzed using up to 2TB ram but greater resources are required. **D**. Protein Membrane integration in CITE-seq dataset. Spearman protein expression correlations scores obtained on k-nearest neighbouring (k=20) integrated latent spaces for all integration methods and original reference CITE-seq dataset k-nearest neighbouring (k=20).

### Sensitivity analysis

Next, we evaluated the robustness of both Seurat4^23^ and LIBRA by reducing the number of cells. Here we randomly selected and removed a certain percentage of cells while computing PPJI for each case. As expected, a decrease in the number of cells diminished Seurat4^23^ and LIBRA’s performance (Fig.3B). Interestingly, when reducing the number of cells, LIBRA performs significantly better than Seurat4^23^. Additional robustness analysis shows LIBRA can maintain high accuracy against randomization of the pairing information, dropout, and overtraining (see Table S3).

### Generalization of the results

To assess the generalizability of LIBRA, we compared LIBRA with the other methodologies using a broader range of datasets. We considered the following datasets: CITE-seq (Human Bone Marrow, DS2^9^), PAIRED-seq (Mouse Adult Cerebral Cortex, DS3^7^) and SHARE-seq (Mouse Skin, DS4^6^), 10X (PBMC, DS5^8^), 10XMultiome (Human Bone Marrow, DS6^28^), CITE-seq (Human Bone Marrow, DS7^28^) and scNMT-seq (Mouse Embryonic Stem Cells, DS8^10^). Further details are provided in Fig.3C, Table S4. PPJI based-comparison was not feasible in DS8 because of the limited number of cells and (as a result) the very limited number of clusters identified. A general observation, see Fig.3C, is that fine-tuned Seurat4^23^ surpasses all other methodologies in most cases. However, Seurat4^23^, MOFA+^22^, and LIBRA are comparable, and depending on the dataset, a different method performs better. BABEL^24^ provides the worse results except for DS3 when compared against ATAC-seq as *ds1*. Interestingly, we found that DS3 ATAC-seq provides limited information on clusters, which is observed on the integration-based clustering from Seurat4^23^ and LIBRA any additional information to the multi-omics integration. However, BABEL^24^ appears to prioritize the information from ATAC-seq in the integration as shown in Fig.S3. It is relevant to note that BABEL^24^ development aimed at prediction, not cell-type identification. We also observed that the normalization procedure (e.g., using or not SCT) has a limited effect on the PPJI analysis (see Table S5). In the case of DS5, being the set with the largest number of cells (at the moment this analysis was carried out), we observed that using “all features’’ instead of “most-variables features” provided slightly better results in the integration (<0.02 PPJI difference); as a result, we analyzed all methods with “all features” option. It was not possible to run BABEL^24^ within a reasonable amount of time (e.g., less than a week) with the entire set of features on DS5.

As an extension to the current work, we have compared against the winner algorithm in the NeurIPS challenge (*concatenated AE*) on dataset DS6, obtaining a resolution of 23 clusters with a PPJI score of 0.72 and 0.64 for scRNA and scATAC, respectively. Considering LIBRA performance with default hyperparameters we obtained a resolution of 28 clusters and PPJI scores of 0.79 and 0.67. We conclude that LIBRA outperforms the concatenated AE in resolution and biological information preservation in SLS.

To evaluate LIBRA in other combinations of data modalities, we investigated the prediction in CITE-seq. To that end, we estimated the expression of 25 protein values in CITE-seq DS2 dataset^8^ using the profiles from the neighbouring cells as conducted in Seurat4^23^ analysis^13^ using previously explained metric computed at each of the SLS components obtained. Seurat4^23^, LIBRA, totalVI^34^, and MOFA+^22^ returned the best results for 14, 11, 1, and 1 of the 25 antibodies, respectively as shown in Fig.3D. Overall, Seurat4^23^ provides more stable results, followed by LIBRA.

### Predictive power of LIBRA

While LIBRA is comparable with Seurat4^23^ using PPJI, the LIBRA framework’s added value is its use as a predictive model. Generating a LIBRA model for a paired dataset, enables the prediction of profiles from single-omic single-cell data of the same biological system. Considering the dt1=ATAC and dt2=RNA, we quantified the predictive power for RNA profiles, *predRNA*, as the Pearson correlation between known and predicted profiles as used in BABEL^24^. We acknowledge that *predRNA* can also be considered as an evaluation measure for NN1. We compared *predRNA* value between BABEL^24^ and LIBRA on all datasets that it was possible (Table S6); we observed that LIBRA outperforms BABEL^24^ in all cases. We also observed that the prediction values estimation is valid for all clusters, and the correlation per cluster is not associated with the number of cells in the cluster (Tables S7,8). Similarly, we also observed that LIBRA outperforms BABEL^24^ when using RNA to predict ATAC-seq profiles (0.87 vs. 0.85 *predATAC*, see Supplementary Materials and Methods).

### Comparing running times

When comparing running times (Table 1), Seurat4^23^ is the fastest. However, because the training of LIBRA is to be performed once or, as a maximum, a few times in any single-cell multi-omics analysis, we consider that the time cost on CPU observed to be functional (Table1). Notably, while BABEL^24^ and LIBRA are both AE-inspired methodologies, the more complex BABEL^24^ architecture makes it significantly more time-consuming.

**Table 1.** Computational costs of each of the methodologies. Times required to generate the integrated spaces for each of the tested models. Estimation for the running on GPUs (Tesla V100) or CPU based systems was carried based on Nvidia supplier specifications.

| DataSet | ID | LIBRA(CPU) | Training time<br>BABEL(GPU) | BABEL GPU 2 CPU | LIBRA(CPU) | Per epoch<br>BABEL(GPU) | BABEL GPU 2 CPU | MOFA+(CPU) | totalVI(CPU) | Seurat4(CPU) |
| --- | --- | --- | --- | --- | --- | --- | --- | --- | --- | --- |
| DS1 | GSE126074 | 12min | 10mins | 6 hours | 3s | 12s | 360s | 6:30 hours | 15 hours | < 3mins |
| DS2 | GSE128639 | 10mins | (*) |  | 1s | (*) |  | 3:42hours | 12 hours | < 3mins |
| DS3 | GSE130399 | 10mins | 5mins | 3 hours | 5s | 8s | 240s | 54min | 4:30 hours | < 3mins |
| DS4 | GSE140203 | 33mins | 23mins | 11:30 hours | 10s | 17s | 128s | 2:24hours | 11:30 hours | < 3mins |
| DS5 | 10X Genom | 25mins(*) | (**) |  | 3s | (**) |  | 3:30 hours | 13 hours | < 3mins |
| DS6 | GSE194122 | 42mins | (*) |  | 10s | 17s | 128s | Out of mem (2TB ram) | > 24 hours | < 3mins |
| DS7 | GSE194122 | 10mins | (*) |  | 3s | (*) |  | 5:30 hours | 19 hours | < 3mins |
| DS8 | GSE109262 | 1mins | (***) |  | 1s | (***) |  | -- | -- | < 3mins |

### Adaptative LIBRA: automatic dataset specific auto-finetuning for LIBRA

LIBRA has the best results in prediction across data modalities. Yet, the default version using the same scheme for any dataset has shown slightly weaker results than Seurat4^23^ for integrative tasks. Based on an observation derived from the NeuRIPS challenge, fine-tuning appears to a necessary step for finding an optimal performance of these neural network architectures. To further investigate such a hypothesis, we computed the evaluation scores of LIBRA for different combinations of parameters while setting the other parameters to a given value using the largest dataset DS6. As shown in Fig.4A,B, the different metrics used may differ in the result for different combinations. Therefore, it is necessary to identify the fitness landscape shown in Fig.4C. In the example, aimed to optimize “*integrative scores*” we have compared 423 models using a grid of vectors for the different hyperparameters (Table S9; Materials and Methods). As a result, see Fig.4D, fine-tuned LIBRA (aLIBRA) outperforms Seurat4^23^, while increasing the PPJI scores for RNA (from 0.79 to 0.82) and ATAC (from 0.67 to 0.81) compared to those obtained by Seurat4^23^ (0.84 and 0.77). See also the extended clustering definition in Fig 4E,F.

**Figure 4.**
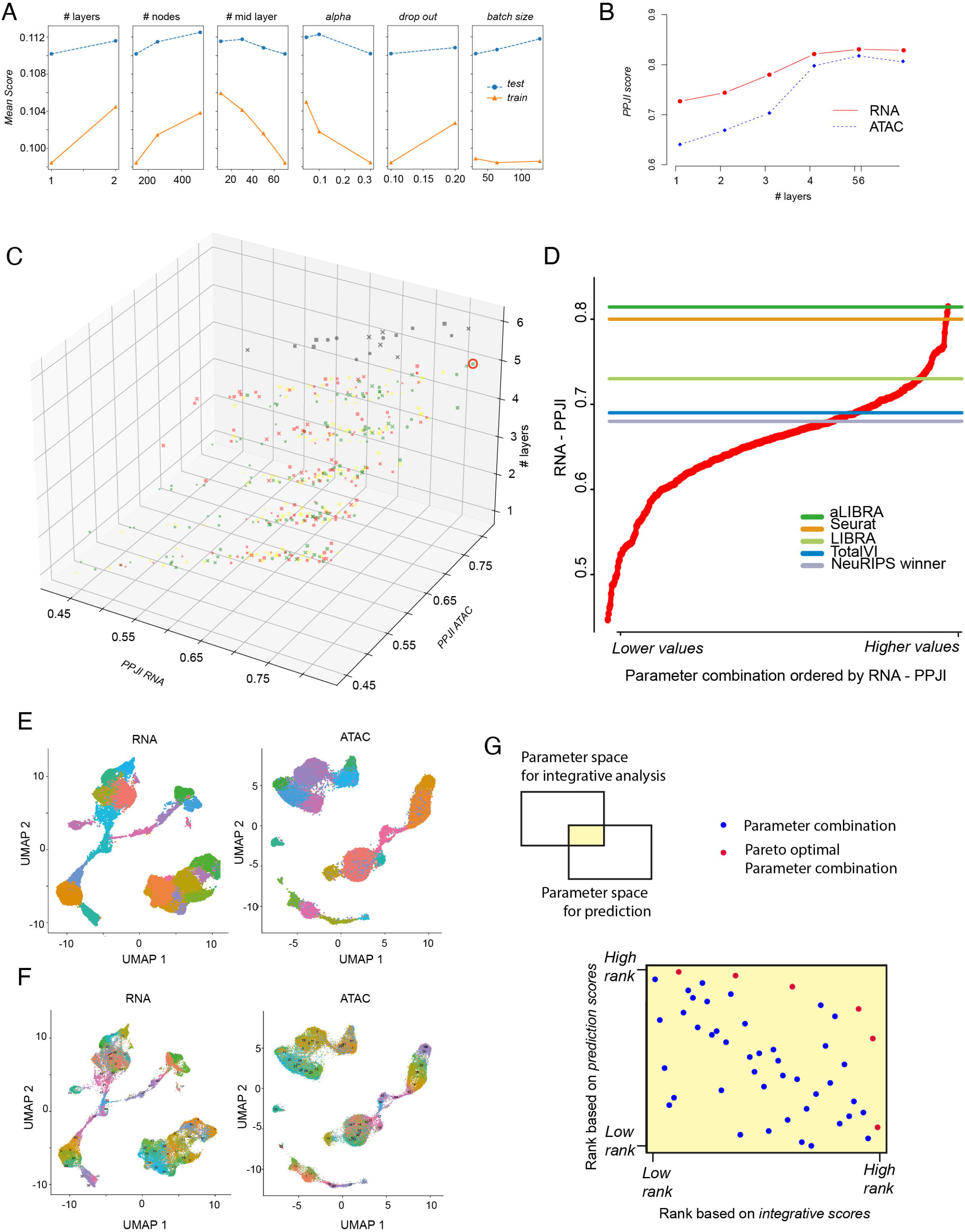
Additional evaluation. **A**. Prediction fine-tuned LIBRA outcomes from hyperparameters tested against MSE metric. Orange represents the performance of model over training set 0.80 of total DS6 and blue the performance obtained over remaining 0.20 test set. **B**. Integration fine-tuned LIBRA outcomes from hyperparameters tested against PPJI metric. Best models for different number of layers are represented showing where plateau is obtained. RNA preserved information score as red and ATAC preserved information score as blue. **C**. aLIBRA results over DS6 (10XMultiome) from NeurIPS challenge scRNA and scATAC PPJI values. The 6D representation provide the differences in performance because of hyperparameter differences. The impact of an adaptative version to surpass fixed configurations limitations provide substantial performance improvements compared to fixed configuration. Surrounded by a red circle the model that has shown a higher performance over the rest combinations of hyperparameters. Each dimension is detonated as; X-axis (rna-seq ppji), Y-axis (atac-seq ppji), Z-axis (# of layers), size (%dropout, 0.1-smaller and 0.2-bigger), colors (#of nodes of first layer, 256-red, 512-yellow, 1024-green and 2048-black) and shape (#of nodes in middle layer, 10-comma, 50-cross and 70-circle). **D**. Ranking of the parameter combinations from lower to higher values for the combinations investigated in the integrative analysis. Lines denote the values obtained by different methodologies. **E**. Original RNA and ATAC clustering information is shown within UMAP representation. **F**. As (D) but information provided corresponds to LIBRA fine-tuned model clustering outcomes. **G**. Upper left: shows that for each task (integration and prediction), two different sets of parameter combinations are investigated. However, there is an overlap. For the overlap, the ranking of both evaluation scores are shown in order to identify the *Pareto optimals*.

Importantly, the parameter space can be defined differently for prediction and for integration. In our case, for instance, we observed that frameworks with lower nodes return better predictions; see in Methods for the details of the two different parameter sets. We investigated if there were combinations – from the overlapping parameter space – that returned good evaluations in both criteria; see Fig4.G. We observed that there are *Pareto optimals*: competitive frameworks in both tasks.

Based on all those observations, we conclude that a predefined or even a step-wise model concept can be further improved when augmented by a fine-tuning strategy that adapts to the data (aLIBRA).

### LIBRA as a resource

LIBRA has been implemented as a Python package called sc-libra. It provides state-of-art performance while retaining good execution timing. In contrast to other AI methods that require longer CPU time to accomplish similar training processes. Furthermore, LIBRA includes the possibility to train hundreds of models in parallel for the fine-tuning of the parameters in a data-driven manner (aLIBRA).

LIBRA is a modular toolbox and is hopefully easy to use in our experience. All outputs from functions and directories tree are generated “*behind the scenes*,” and the required user interaction is very limited Fig.5(A).

**Figure 5.**
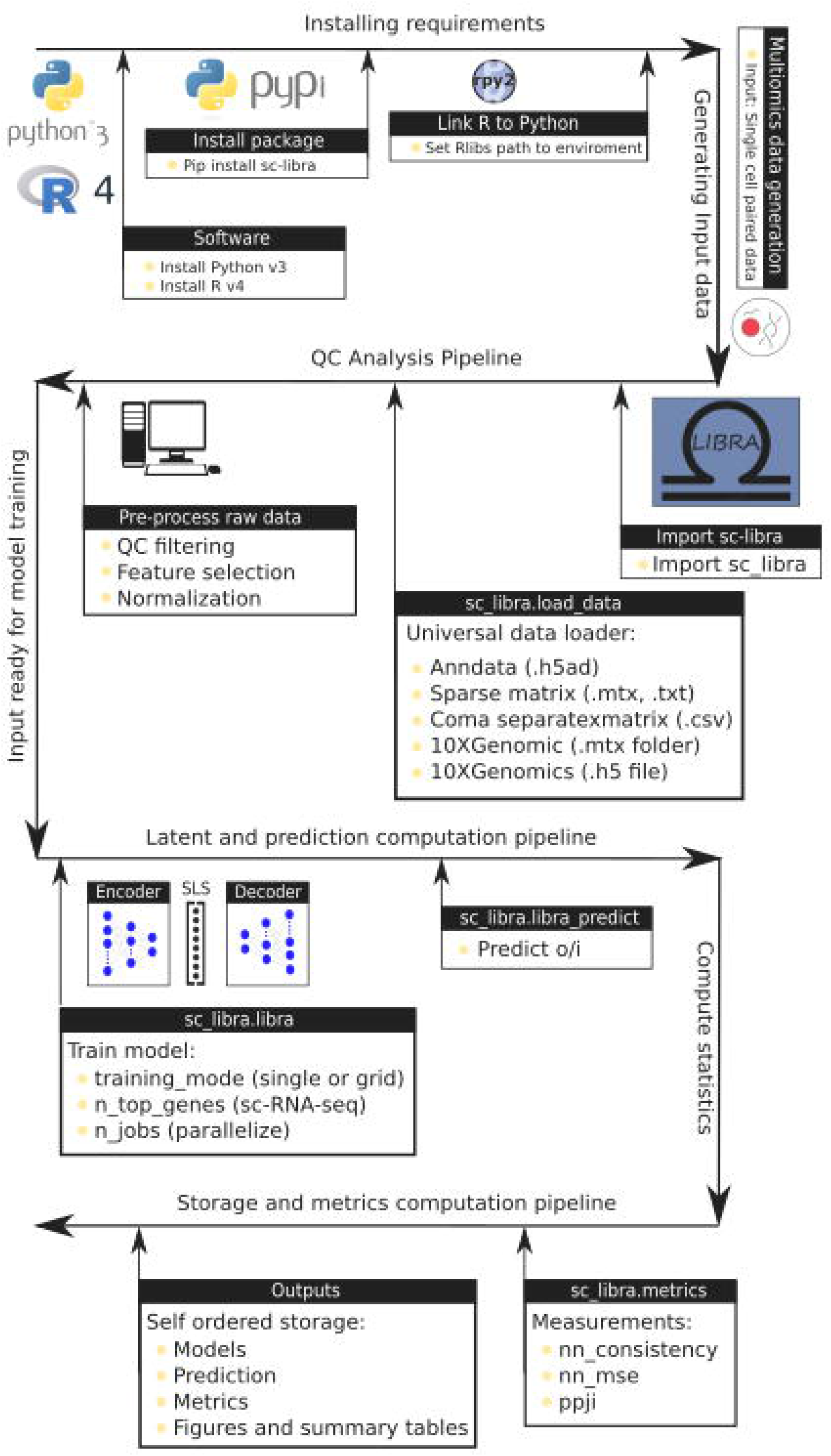
Libra as users resource. sc-libra Python package graphical pipeline. From top to bottom its represented the guide to properly get and run LIBRA pipeline with expected outputs and metrics.

## Discussion

The deal with the avalanche of emerging multi-omics profiling technologies^35^, the community needs powerful computational tools for multi-omics data analysis^11^. Such a synergistic development holds promise for deep deconvolution^36^ and predictive modelling of cellular layers of genomic circuits for health and aberrations underpinning diseases^37^. Furthermore, those tools must adapt to different data modalities and each dataset’s specific characteristics. To meet this demand, we introduced LIBRA.

LIBRA is a tool that leverages paired-single-cell information using an AE framework to address two fundamental challenges when analyzing multi-modal single-cell data. Namely, identifying the joined space, therefore facilitating cell-type resolution and enabling prediction between different omics modalities. We observed that LIBRA compares competitively with state-of-the-art tools in both tasks and is robust when the number of cells is reduced. Furthermore, the LIBRA architecture and learning scheme is generalizable to any pair of omics.

The limited CPU time of the model allows its adaptation to different datasets (with its considerable effect on performance enhancement), something that would be impossible with tools that require larger CPU times such as totalVI^34^, MOFA+^22,^ or BABEL^24^. Additionally, LIBRA model’s simplicity, limited CPU time requirements, and scalability allow it to be combined with a fine-tuning strategy. The *aLIBRA* finetunes and improves the outcome of the model significantly and consequently outperforms other methodologies. The improvement observed in aLIBRA when compared with LIBRA aligns with the observation from NeurIPS challenge that fine-tuning was required for top-scoring methodologies. Furthermore, identifying frameworks during the fine-tuning that are competitive in both tasks (prediction and integration) is feasible as shown in Fig.4G.

In summary, LIBRA and aLIBRA are state-of-the-art tools for multi-modal single-cell analysis for *prediction* and *projection*, whose implementations are available as OpenSource in R and Python, with tutorials available. LIBRA is implemented as a Python package (under PyPI repository) named sc-libra, allowing users to efficiently perform all the proposed analyses and metrics on any pair of paired single cell omics. Online docs for sc-libra are provided for user guide through this package.

### Availability and requirements

Project name: LIBRA

Project home page: https://github.com/TranslationalBioinformaticsUnit/LIBRA.

Operating system(s): Platform independent. Tested on LINUX.

Programming language(s): sc-libra (LIBRA package implementation at PyPI), Python, Jupyter notebook, R and RMarkDown.

License: GPL-3.0 License

Any restrictions to use by non-academics: None

### Availability of data and materials

The datasets re-analysed during the current study are available in the NCBI GEO repository via accession numbers GSE126074, GSE128639, GSE130399, GSE140203, GSE194122, GSE109262 and 10X Genomics website repository. The developed package and it’s online documentation and the code used for the re-analysis, are available at:

sc-libra package:

https://pypi.org/project/sc-libra/

sc-libra online docs:

https://sc-libra.readthedocs.io/en/latest/

GitHub repository:

https://github.com/TranslationalBioinformaticsUnit/LIBRA

Cone of GitHub repository plus data repository:

https://figshare.com/articles/journal_contribution/LIBRA-main_zip/19466246

## Abbreviations

NN: Neural networks
GEO: Gene Expression Omnibus
SLS: Shared latent space
PJI: Pairwise Jaccard Index
DS: Data set
predRNA: Predicted RNA
predATAC: Predicted ATAC
MSE: Mean squared error
SNARE-seq: Droplet based technology to profile chromatin accessibility and gene expression from the same cells.
CITE-seq: Qualitative information over gene expression and surface proteins with available antibodies on a single cell level.
Paired-seq: Combinatorial indexing strategy to simultaneously tag both the open chromatin fragments generated by the Tn5 transposases and the cDNA molecules generated from reverse transcription.
SHARE-seq: Strategy that uses three rounds of barcodes by ligating barcoded adaptors to both RNA (gene expression) and tagmented DNA (chromatin accessibility) to achieve the multi-omic profiling from the same single cells.
10X: 10X Genomics *Single*-*Cell* Multiomics Solutions
CITE-seq: Method for performing RNA sequencing along with gaining quantitative and qualitative information on surface proteins with available antibodies on a single cell level.
scNMT-seq: Method to look at methylation (CpG) and chromatin accessibility (GpC).

## Acknowledgements

Not applicable.

## Author information

### Affiliations

Navarrabiomed, Complejo Hospitalario de Navarra (CHN), Universidad Pública de Navarra, IdiSNA, Pamplona, Spain, 31008.

Xabier Martinez-de-Morentin, David Gomez-Cabrero

Biological and Enviromental Science and Engineering Division, King Abdullah University of Sience and Technology, Thuwal, Saudi Arabia, 23955-6900.

Sisi Qu, Sumeer A. Khan, Robert Lehmann, Alberto Maillo, Jesper Tegner, David Gomez-Cabrero

Department of Oncology and Pathology Cancer Center Karolinska, Karolinska Institute, Stockholm, Sweden, 171 77.

Narsis Kiani

Hematology-Oncology Program, Center for Applied Medical Research (CIMA), University of Navarra, Instituto de Investigación Sanitaria de Navarra (IdiSNA), Navarra, Spain, 31008.

Felipe Prosper

Service of Hematology and Cell Therapy, Clínica Universidad de Navarra, Pamplona, Spain, 31008.

Felipe Prosper

Computer, Electrical and Mathematical Sciences and Engineering Division, King Abdullah University of Science and Technology, Thuwal, Saudi Arabia, 23955-6900.

Jesper Tegner

Centre for Host Microbiome Interactions, Faculty of Dentistry, Oral & Craniofacial Sciences, King’s College, London, UK, 7836 5454.

David Gomez-Cabrero

## Contributions

XMM, DGC designed LIBRA and the computational experiments shown. XMM performed most of the experiments and analysed the results. SQ conducted the experiments associated with BABEL. XMM, JT and DGC wrote the first draft and the final version. SQ, SK, RL, AM, NK, FP, and JT, provided additional insights into the experiments and the text. All authors reviewed the manuscript before submission.

## Ethics declarations

## Ethics approval and consent to participate

Not applicable.

## Consent for publication

Not applicable.

## Competing interests

The authors declare no competing interests.

## Additional information

Not applicable.

## Supplementary Information

Additional file 1

Supplementary Information: Supplementary Materials and Methods, Github repository, Supplementary figures, Supplementary tables and references.

## Rights and permissions

## Notes

### Competing Interest Statement

The authors have declared no competing interest.

### Summary of Updates

- Analysis considering additional data-sets. - Includes analysis of the computational time requirements. - Including automatic fine-tuning.

## References

1. Pratapa, Aditya; Jalihal, Amogh P.; Law, Jeffrey N.; Bharadwaj, Aditya; Murali, T. M. (2020). Benchmarking algorithms for gene regulatory network inference from single-cell transcriptomic data. Nature Methods 17, 147–154 (2020).

2. Tan, K., Tegner, J. & Ravasi, T. Integrated approaches to uncovering transcription regulatory networks in mammalian cells. Genomics 91, 219–231 (2008).

3. Schier, A. F. Single-cell biology: beyond the sum of its parts. Nat. Methods 17, 17–20 (2020).

4. Chen, S., Lake, B. B. & Zhang, K. High-throughput sequencing of the transcriptome and chromatin accessibility in the same cell. Nat. Biotechnol. 37, 1452–1457 (2019).

5. Cao, J. et al. Joint profiling of chromatin accessibility and gene expression in thousands of single cells. Science (80-.). 361, 1380–1385 (2018).

6. Ma, S. et al. Chromatin Potential Identified by Shared Single-Cell Profiling of RNA and Chromatin. Cell 183, 1103–1116.e20 (2020).

7. Zhu, C. et al. An ultra high-throughput method for single-cell joint analysis of open chromatin and transcriptome. Nat. Struct. Mol. Biol. 26, 1063–1070 (2019).

8. Stuart, Tim; Butler, Andrew; Hoffman, Paul; Hafemeister, Christoph; Papalexi, Efthymia; Mauck, William M.; Hao, Yuhan; Stoeckius, Marlon; Smibert, Peter; Satija, Rahul. Comprehensive Integration of Single-Cell Data. Cell. 177, 1888–1902 (2019).

9. Stoeckius, M. et al. Simultaneous epitope and transcriptome measurement in single cells. Nat. Methods 14, 865–868 (2017).

10. Clark, S. J. et al. scNMT-seq enables joint profiling of chromatin accessibility DNA methylation and transcription in single cells. Nat. Commun. 9, 781 (2018).

11. Argelaguet, R., Cuomo, A.S.E., Stegle, O. et al. Computational principles and challenges in single-cell data integration. Nat Biotechnol 39, 1202–1215 (2021).

12. Rohart, F., Gautier, B., Singh, A., & Lê Cao, K.-A. mixOmics: An R package for ‘omics feature selection and multiple data integration. PLOS Computational Biology, 13, (2017).

13. Argelaguet, Ricard; Velten, Britta; Arnol, Damien; Dietrich, Sascha; Zenz, Thorsten; Marioni, John C; Buettner, Florian; Huber, Wolfgang; Stegle, Oliver. Multi-Omics Factor Analysis—a framework for unsupervised integration of multiomics data sets. Molecular Systems Biology, 14, (2018).

14. Lock, E. F., Hoadley, K. A., Marron, J. S., & Nobel, A. B. Joint and individual variation explained (JIVE) for integrated analysis of multiple data types. The Annals of Applied Statistics, 7, (2013).

15. Teschendorff, A. E., Jing, H., Paul, D. S., Virta, J., & Nordhausen, K. Tensorial blind source separation for improved analysis of multi-omic data. Genome Biology, 19, (2018).

16. Gomez-Cabrero, David; Tarazona, Sonia; Ferreirós-Vidal, Isabel; Ramirez, Ricardo N.; Company, Carlos; Schmidt, Andreas; Reijmers, Theo; Paul, Veronica von Saint; Marabita, Francesco; Rodríguez-Ubreva, Javier; Garcia-Gomez, Antonio; Carroll, Thomas; Cooper, Lee; Liang, Ziwei; Dharmalingam, Gopuraja; van der Kloet, Frans; Harms, Amy C.; Balzano-Nogueira, Leandro; Lagani, Vincenzo; Tsamardinos, Ioannis; Lappe, Michael; Maier, Dieter; Westerhuis, Johan A.; Hankemeier, Thomas; Imhof, Axel; Ballestar, Esteban; Mortazavi, Ali; Merkenschlager, Matthias; Tegner, Jesper; Conesa, Ana. STATegra, a comprehensive multi-omics dataset of B-cell differentiation in mouse. Scientific Data, 6, 256– (2019).

17. Stegle, Oliver; Teichmann, Sarah A.; Marioni, John C. Computational and analytical challenges in single-cell transcriptomics. Nature Reviews Genetics, 16, 133–145 (2015).

18. Jeffrey M. Perkel. Single-cell analysis enters the multiomics age. Nature (2021).

19. Marx, V. How single-cell multi-omics builds relationships. Nat Methods 19, 142–146 (2022).

20. Stuart, T. et al. Comprehensive Integration of Single-Cell Data. Cell 177, 1888–1902 (2019).

21. Argelaguet, R., Cuomo, A. S. E., Stegle, O. & Marioni, J. C. Computational principles and challenges in single-cell data integration 39, 1202–1215 Nat. Biotechnol (2021).

22. Argelaguet, R. et al. MOFA+: a statistical framework for comprehensive integration of multi-modal single-cell data. Genome Biol. 21, 111 (2020).

23. Hao Y, Hao S, Andersen-Nissen E, Mauck WM 3rd, Zheng S, Butler A, Lee MJ, Wilk AJ, Darby C, Zager M, Hoffman P, Stoeckius M, Papalexi E, Mimitou EP, Jain J, Srivastava A, Stuart T, Fleming LM, Yeung B, Rogers AJ, McElrath JM, Blish CA, Gottardo R, Smibert P, Satija R. Integrated analysis of multi-modal single-cell data. Cell. 184, 3573–3587 (2021).

24. Wu, K. E., Yost, K. E., Chang, H. Y. & Zou, J. BABEL enables cross-modality translation between multiomic profiles at single-cell resolution. Proc. Natl. Acad. Sci. 118, (2021).

25. Fortelny, Nikolaus; Bock, Christoph. Knowledge-primed neural networks enable biologically interpretable deep learning on single-cell sequencing data. Genome Biology, 21, 190–, (2020).

26. Ravindra, N., Sehanobish, A., Pappalardo, J. L., Hafler, D. A., & van Dijk, D. Disease state prediction from single-cell data using graph attention networks. Proceedings of the ACM Conference on Health, Inference, and Learning, (2020).

27. Kimmel JC, Kelley DR. Semisupervised adversarial neural networks for single-cell classification. Genome Res 31, 1781–1793 (2021).

28. Malte D Luecken, Daniel Bernard Burkhardt, Robrecht Cannoodt, Christopher Lance, Aditi Agrawal, Hananeh Aliee, Ann T Chen, Louise Deconinck, Angela M Detweiler, Alejandro A Granados, Shelly Huynh, Laura Isacco, Yang Joon Kim, Dominik Klein, BONY De Kumar, Sunil Kuppasani, Heiko Lickert, Aaron McGeever, Honey Mekonen, Joaquin Caceres Melgarejo, Maurizio Morri, Michaela Müller, Norma Neff, Sheryl Paul, Bastian Rieck, Kaylie Schneider, Scott Steelman, Michael Sterr, Daniel J. Treacy, Alexander Tong, Alexandra-Chloe Villani, Guilin Wang, Jia Yan, Ce Zhang, Angela Oliveira Pisco, Smita Krishnaswamy, Fabian J Theis, Jonathan M. Bloom. A sandbox for prediction and integration of DNA, RNA, and proteins in single cells. OpenReview.net (2021).

29. Wolpert, D.H., Macready, W.G. (1997), “No Free Lunch Theorems for Optimization”, IEEE Transactions on Evolutionary Computation 1, 67.

30. Kyunghyun, C. et al. Learning Phrase Representations using RNN Encoder-Decoder for Statistical Machine Translation. arXiv (2014).

31. Pedregosa, F. and Varoquaux, G. and Gramfort, A. and Michel, V. and Thirion, B. and Grisel, O. and Blondel, M. and Prettenhofer, P. and Weiss, R. and Dubourg, V. and Vanderplas, J. and Passos, A. and Cournapeau, D. and Brucher, M. and Perrot, M. and Duchesnay, E. Titulo. Journal of Machine Learning Research 12, 2825–2830 (2011).

32. Bing Xu, Naiyan Wang, Tianqi Chen, Mu Li. Empirical Evaluation of Rectified Activations in Convolutional Network. arXiv(2015).

33. Sammut C., Webb G.I. Mean Squared Error. Encyclopedia of Machine Learning. Springer, Boston, MA. (2011).

34. Gayoso, A. et al. Joint probabilistic modeling of single-cell multi-omic data with totalVI. Nat. Methods 18, 272–282 (2021).

35. Mimitou, E. P. et al. Scalable, multi-modal profiling of chromatin accessibility, gene expression and protein levels in single cells. Nat. Biotechnol. (2021) doi:10.1038/s41587-021-00927-2.

36. Zenil, H., Kiani, N. A., Zea, A. A. & Tegnér, J. Causal deconvolution by algorithmic generative models. Nat. Mach. Intell. 1, 58–66 (2019).

37. Tegnér, J. N. et al. Computational disease modeling - fact or fiction? BMC Syst. Biol. 3, 56 (2009).

