## Supplementary figures and images for "Adaptative Machine Translation between paired Single-Cell Multi-Omics Data"

### Fig. S1

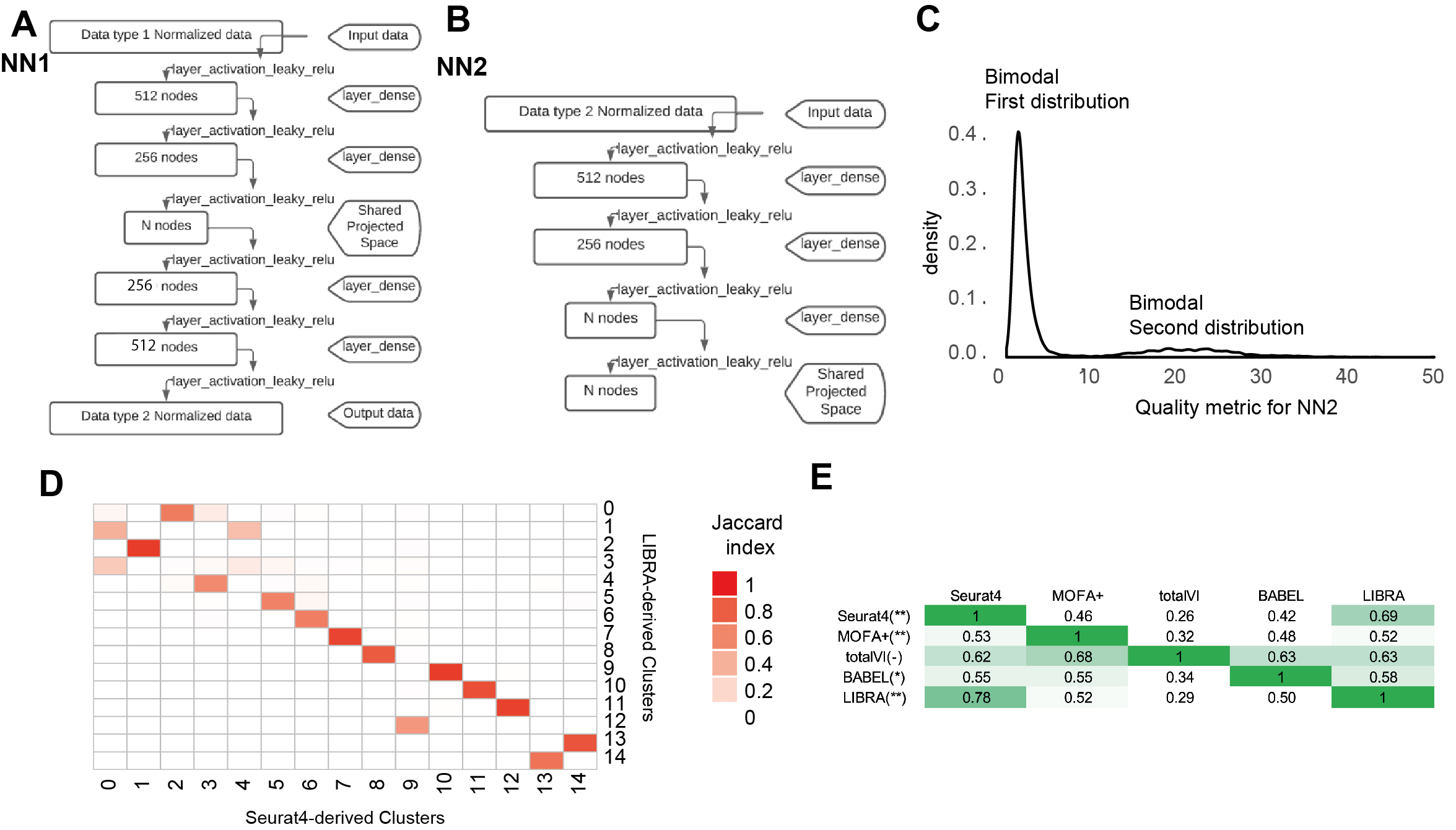

### Fig. S2

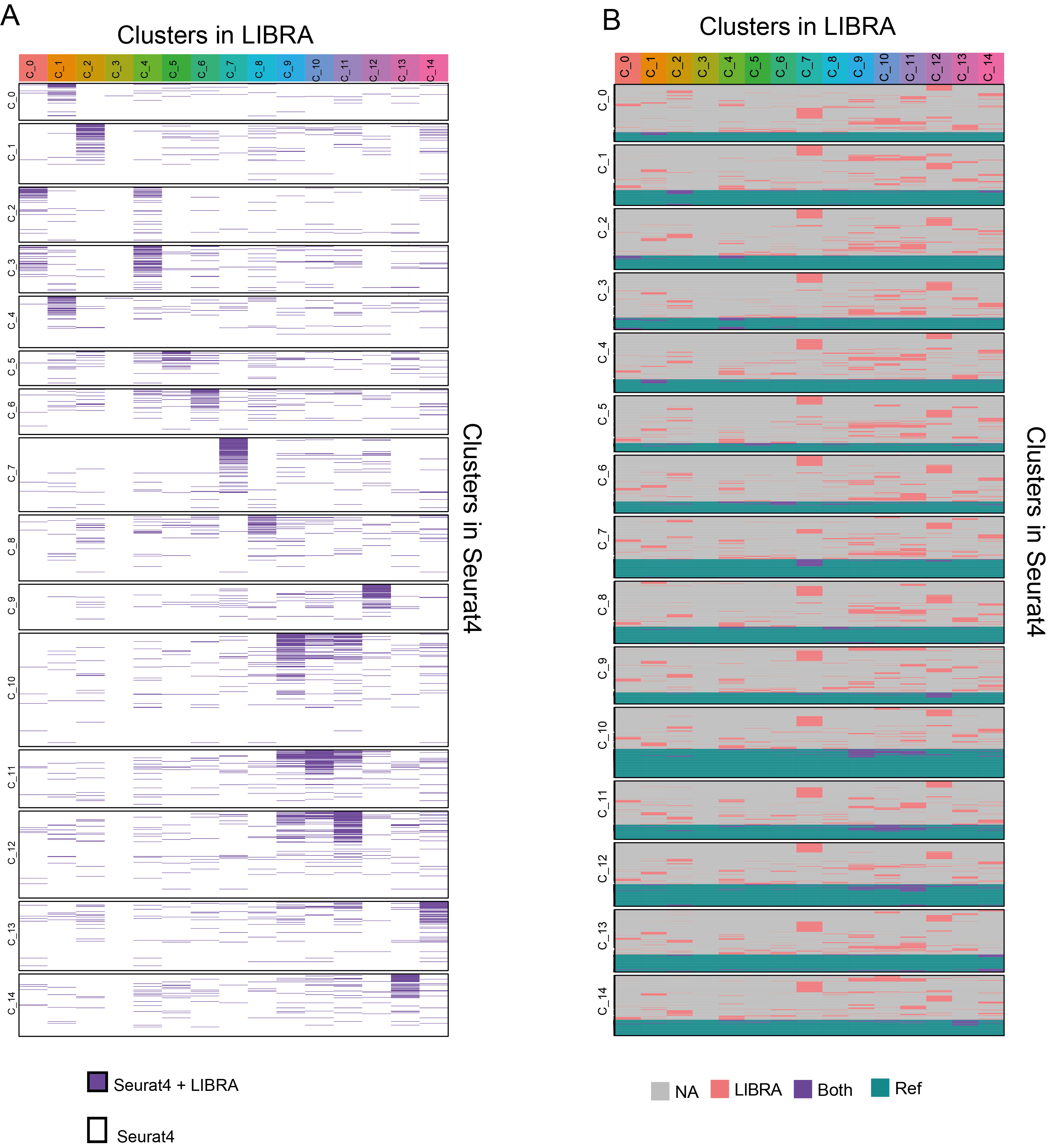

### Fig. S3

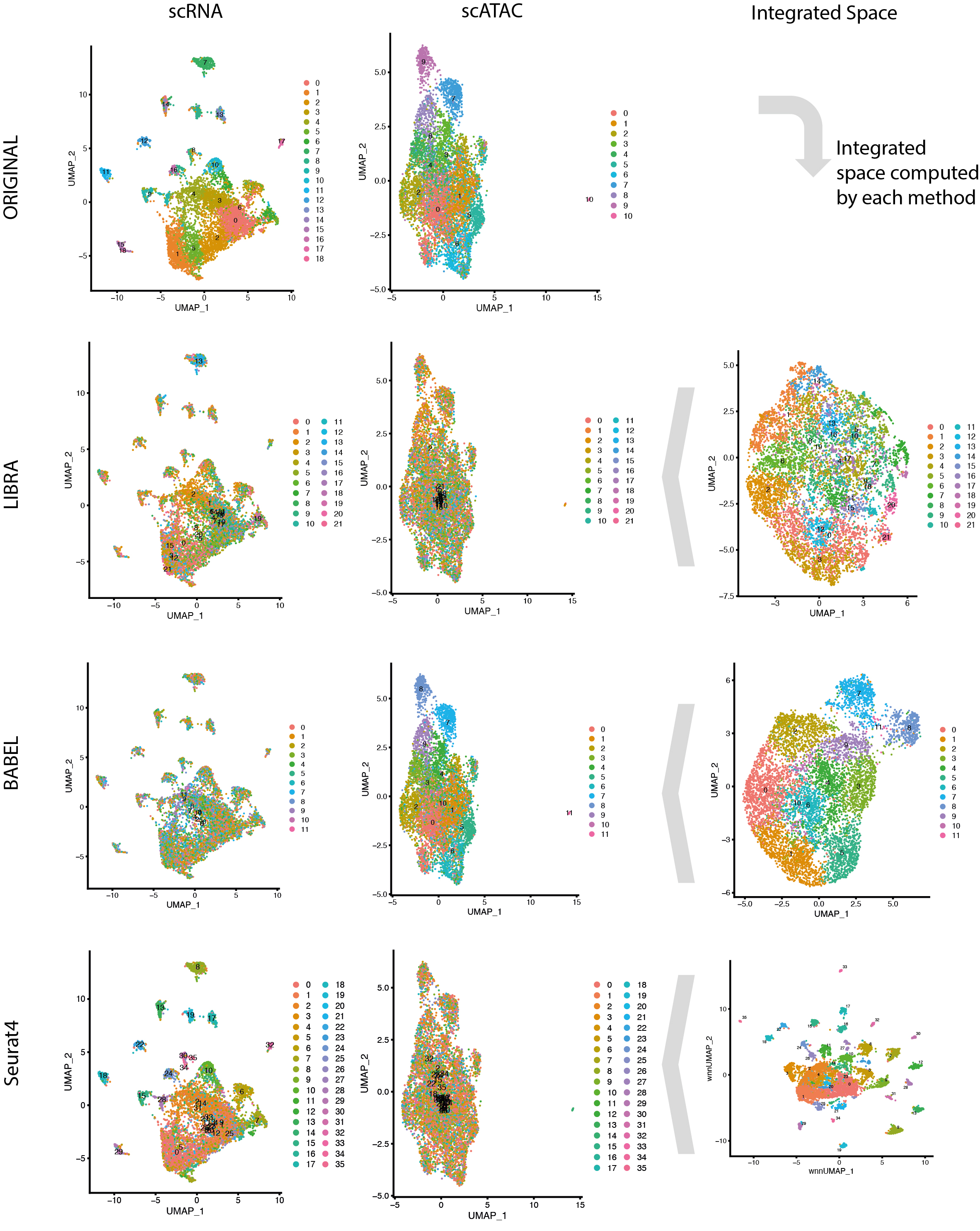
