## Supplementary material for "Adaptative Machine Translation between paired Single-Cell Multi-Omics Data": Supp. Materials

**Supp. Information**


SUPPLEMENTARY METHODS 2

**Datasets.** 2

**Pre-analysis and data processing:** 4

**Visualization:** 5

**Clustering for integrated and independent omic modalities:** 6

**Overview of the Quality Measures.** 6

**Computing the Preserved Pairwise Jaccard Index (PPJI)** 8

**Q1 and Q2 evaluation.** 9

**PredRNA** 9

**PredATAC** 10

**Default model development.** 10

**LIBRA structure and additional hyperparameters.** 11

**LIBRA fine-tune (deep adaptive structure).** 12

GITHUB REPOSITORY 13

SUPPLEMENTARY FIGURE LEGENDS. 14

SUPPLEMENTARY TABLE LEGENDS. 15

REFERENCES 16

### Supplementary Methods

#### *Datasets.*

For developing and testing the performance and quality of LIBRA, eight sets of paired multimodality datasets were used. Additional information is provided in Fig.3C.

DataSet1, SNARE-seq[1]:

- GSE126074.
- Data modalities: single-cell RNA-seq and single-cell ATAC-seq.
- Single-cell nucleus.
- Cells selected: Mouse Adult Cerebral Cortex
- Total number of cells before filtering: 10,309 nuclei.
- Total number of cells after filtering: 6,735 nuclei.

DataSet2, CITE-seq[2,3]:

- GSE128639.
- Data modalities: single-cell RNA-seq and ADT panel for 25 antibodies.
- Cells selected: Human Bone Marrow cells
- Total number of cells before filtering: 33,454 cells.
- Total number of cells after filtering: 30,672 cells.

DataSet3, Paired-seq[4]:

- GSE130399.
- Data modalities: single-cell RNA-seq and single-cell ATAC-seq.
- Cells selected: Mouse Adult Cerebral Cortex
- Total number of cells before filtering: 11,544 cells.
- Total number of cells after filtering: 7,743 cells.

DataSet4, SHARE-seq[5]:

- GSE140203.
- Data modalities: single-cell RNA-seq and single-cell ATAC-seq.
- Single-cell nucleus.
- Cells selected: Mouse Skin
- Total number of cells before filtering: 42,948 cells.
- Total number of cells after filtering: 34,774 cells.

DataSet5, 10X Multiome:

- 10X Genomics website repository.
- Data modalities: 10X Genomics single-cell RNA-seq and single-cell ATAC-seq.
- Single-cell nucleus.
- Cells selected: Human PBMC
- Total number of cells before filtering: 11,709 cells.
- Total number of cells after filtering: 10,412 cells.

DataSet6, 10X Multiome:

- GSE194122
- Data modalities: 10X Genomics single-cell RNA-seq and single-cell ATAC-seq.
- Cells selected: Bone Marrow
- Total number of cells after filtering: 42,492 cells.

DataSet7, CITE-seq:

- GSE194122
- Data modalities: single-cell RNA-seq and ADT panel for 134 antibodies.
- Cells selected: Bone Marrow.
- Total number of cells after filtering: 66,175 cells.

DataSet8, scNMT-seq[6]:

- GSE109262.
- Data modalities: single-cell RNA-seq, single-cell ATAC-seq and single-cell DNA Methylation.
- Single-cell nucleus.
- Cells selected: Mouse Embryonic Stem Cells.
- Total number of cells before filtering: 100 cells.
- Total number of cells after filtering: 69 cells.

Importantly, during the pre-processing and after quality filter the total number of cells used in the analysis were reduced. **The final total numbers of cells used for all the analysis conducted in the paper are described in Table.S4**.

#### *Pre-analysis and data processing:*

DataSet1 (Detailed example, similar pipeline for paired RNA and ATAC data-sets):

*Quality filter – low quality features*: removing low-quality features and cells for both modalities. All cells with an overall abundance level of "number of features per cell" and "number of counts per cell" lower than quantile 0.1 or higher quantile 0.9 were removed from downstream analysis. Minimum abundance filtering was applied for mRNA (ATAC): genes (peaks) profiled in less than 4 cells (3 cells), and cells will lower than 201 genes quantified were filtered. No minimum of peaks per cell was required. After quality and abundance filtering, we considered for the analysis a total of 8,086 cells for scRNA and 8,214 cells for scATAC adult samples.

*ATAC-derived gene activity*: when ATAC-derived gene activity was computed, the Seurat3[7] 'CreateGeneActivityMatrix' function was used with "upstream=2000" bases. And GRCh38 genome as a reference to later identify marker genes over the integrated expression subspace.

*Quality filter – mitochondrial:* 5% mitochondrial filtering was used for the expression matrixes of scRNA and scATAC; in the ATAC case using activity from peaks.

*Variable feature selection*: For variable features selection in scATAC the function 'FindTopFeatures' from Signac[8] library was applied with cut-offs of q0 and q5 (later selected q0 based on higher performance results). The top 2,000 most variable genes (mvg) were selected for scRNA (additionally, 3,000 was tested for mvg with no improvement on performance – data not shown).

*Component parameters*: A total of 15 principal components (PCA) were selected for scRNA reduction, and 50 latent semantic indexing components (LSI) were selected for scATAC.

*The final number of cells*: from the resulting pipeline, a total of 6,735 paired cell profiles were considered for the downstream analysis.

*Integration*: To generate a reference integrated version on this dataset Seurat 3 (unpaired) and Seurat 4 (paired) has been used **over the independent modalities** for further processing and later integration. Using standard normalization and integration guides for both Seurat3 and Seurat4 Weighted Nearest Neighbour Analysis vignette[1]. In Seurat3, *FindTransferAnchors* function using RNA as reference and ATAC as query modalities employing CCA as reduction method was used for anchorset generation, followed by *TransferData* function where anchorset generated was used for transfer into ATAC modality the RNA derived information using LSI dimensional reduction for the weighting anchors. Finally, data was merged

[1] https://satijalab.org/seurat/articles/weighted_nearest_neighbor_analysis.html

for coembed dataset generation. *FindMultimodalNeighbors* function from Seurat4 was used for identifying multimodal neighbors.

Specifics for DataSet2:

- We followed the same pre-processing for the scRNA-seq data analysis as in DataSet1.
- ADT dataset was analysed following Seurat4 Weighted Nearest Neighbour Analysis vignette [same REF as before]. Exact same functions and parameters where used.

Specifics for DataSet6 and DataSet7:

- Data has been used as provided in open problems in Single-Cell Analysis competition (2021). Data is already filtered based on quality controls and normalized using Seurat standard pipeline.

Specifics for DataSet8:

- We followed the same pre-processing for the scRNA-seq and scATAC-seq data analysis as in DataSet1.
- For DNA methylation analysis, first a summarized matrix containing unique CpG positions identified in at least a single-cell as rows and all the cells as columns was generated and filled with obtained percentages of methylation for each corresponding cell and position. Next, DNA methylation values were scaled from 0 to 1, and a binarization was conducted by replacing values greater or equal to 0.5 by 1 and the rest by 0. All positions not methylated in at least 40% of cells were removed based on visual distribution inspection.

*SCT uni-omics integration pre-processing*.

The impact of the normalization during data pre-processing has been tested using a new normalization technique recently proposed by the Seurat team (SCT, sctransform) in DataSet5, PBMC. To accomplish this analysis *SCTransform* function was used with default parameters over RNA dataset, generating the resulting Pearson normalized residuals later used on LIBRA for model training.

#### *Visualization:*

UMAP dimensionality reduction method was applied for visualizations generation over reduced components generated for either independent modalities or integrated modality.

UMAP 2-dimensional reduced space was computed from corresponding low dimensional spaces: RNA(pca), ADT(pca), ATAC(lsi) and DNA methylation(lsi). Implementation used was RunUMAP function from Seurat package.

##### *Clustering for integrated and independent omic modalities:*

All the clustering analysis has been conducted using Seurat Louvain clustering implementation[9]. Depending on the analysis different inputs are considered:

- Single-cell RNA-seq data: PCA components.
- Single-cell ATAC-seq data: LSI components.
- ADT: PCA components.
- Single-cell DNA methylation: LSI components.
- Integrated space: cells were clustered base on shared components generated by the methods considered (LIBRA, Seurat4, MOFA+[10], totalVI[11] and BABEL[12]). The number of components used over these shared spaces were 10, 15, 15, 10 and 16 respectively.

Louvain resolution has been fixed to default 0.8 value for integrated subspaces while the starting seed has been computed over 1,000 times for excluding spurious additional clusters that can appear because of an arbitrary starting seed selection for Louvain clustering. The number of nearest neighbours has been used as default K=30 but optimized using a *snakemake workflow*[2] which is based on subsampling and repeated clustering using *scclusteval*[13] as a reference. The bootstrap related parameters used where: subsample rate of 0.8 for a total of 20 subsamples using values of k from 8 to 16 (in steps of +2) and subsample resolutions from 0.6 to 1.4 (in steps of +0.2) generating a total of 500 analysis.

##### *Overview of the Quality Measures.*

To evaluate the quality of LIBRA, we designed several quality metrics. The first metric measures the quality of the first NN (**referred to as Q1, and computed as MSE**), while the second, **Q2**, measures the accuracy of the dt2 projection (Fig.2B) using an euclidean distance (which is the square root of the MSE used during training). When evaluating Q2 per-cell, we identified a bimodal distribution of the euclidean estimated distances per cell; such distribution was associated to whether the cells were part of the training or not (Fig.S1C); thus, both distributions of the bimodal were evaluated separately.

To evaluate the additional value within the shared latent space (SLS), we designed the **Preserved Pairwise Jaccard Index** (PPJI), a non-symmetric distance metric between

---

[2] https://github.com/crazyhottommy/pyflow_seurat_parameter

two clusterings. PPJI compares clusters derived from single-omics and multi-omics profiling (Fig.2B). For instance, when comparing RNA derived clustering (A) and CPS derived clusterings (B), the PPJI computes the Jaccard Index between every pair of clusters from A and B and summarizes the value (e.g., sum) over the clusters of B (Fig.2C). Unless the two clusterings have the same number of clusters, the outcome will not be symmetric. PPJI is aimed to measure how well the CPS projection provides a finer granularity than a single-omics.

Using these three quality measurements (Q1, Q2, PPJI) and the SNARE-seq[1] adult brain mouse dataset, we selected the hyperparameters for LIBRA (see details in the "Default Model Development Section" and Table.S1):

(i) Autoencoder-type configuration,
(ii) The number of dimensions of the projected space,
(iii) Selection between peak and gene-activity derived information for ATAC-seq,
(iv) The assignation of dt1 and dt2,
(v) The features to be included (all vs most variable features (MVF)) and
(vi) The number of intermediate layers.

In all cases, we favoured first maximal PPJI values, then minimal Q2, and finally, minimal Q3 values. The LIBRA' selected parameters were: *AE-based framework, ten dimensions for the middle layer, dt1=ATAC > dt2=RNA, MVP features included, and two hidden layers* (see details in the "Default Model Development Section").

#### *Computing the Preserved Pairwise Jaccard Index (PPJI)*

Given a set of cells S and two clustering's A and B of the cells, Jaccard Similarity Index (JSI) is computed as follows for every pair of clusters i ∈ A and $j$ ∈ B ($a_i$ and b*j* denote respectively the set of cells in cluster *i* and *j*):

$$JSI(a_i, b_j) = \frac{(a_i \cap b_j)}{(a_i \cup b_j)}$$

The function *PairWiseJaccardSetsHeatmap* from scclusteval[13] package was used for Jaccard Similarity Index estimation. The Pairwise Jaccard Index is a matrix that computes JSI for all pairs of clusters *i* ∈ A and j ∈ B.

When we aim to investigate how A clustering projects on B clustering, by using the matrix, for each cluster i ∈ A, the following statistic is computed:

$$S_i = \sum_{j \in B} JSI(a_i, b_j)$$

Finally, PPJI is computed as follows, where *I* is the total number of clusters in A:

$$\mathbf{PPIJ} = \frac{\mathbf{1}}{\boldsymbol{I}} \sum \boldsymbol{S}_{\mathbf{i}}$$

It is important to note that PPJI is not a symmetric measure, especially in the case where the number of clusters is different.

##### *Q1 and Q2 evaluation.*

Mean square error (MSE) is used as a loss function in the training of networks NN1 and NN2. For instance, in the case of NN1:

$$Q1 = MSE_{NN1} = \sum_{i=1}^{n} (dt_{2,i} - \widehat{dt}_{2,i})^2$$

Where $\widehat{dt}$ denotes the estimated value and *dt* denotes the original value. n is the total number of cells. Q1 is reported as two values:

- Q1 training, the MSE over the training set of cells, when computed for all cells.
- Q1 test, the MSE value when applied to all test cells. Test cells are a 20% of the original number of cells.

And, while NN2 is trained using MSE as a loss function, the Q2 returned is the root square of MSE when computed for all cells, which effectively is the Euclidian distance between predicted value and expected value.

$$Q2 = \sqrt[2]{MSE_{NN2}}$$

Q2 is computed separately for two different set of cells, due to the observation that Q2 follows a bimodal distribution (Fig.1S(c)) an statistic are computed for each of the two distributions identified in the bimodal.

##### *PredRNA*

RNA prediction was obtained by first loading hdf5 format stored trained LIBRA model from ATAC(dt1) to RNA(dt2), then *predict* Keras function was used for getting output from output layer (RNA dt2 like) from ATAC(dt1) input data as parameters.

Evaluation for quality check was performed by computing Pearson correlation between each pair of cells from predicted RNA and original RNA training input data. This computation was performed using *cor* function from *stats* package.

#### *PredATAC*

ATAC prediction was obtained by first loading hdf5 format stored trained LIBRA model from RNA(dt1) to ATAC(dt2), then *predict* Keras function was used for getting output from output layer (ATAC dt2 like) from RNA(dt1) input data as parameters.

Evaluation for quality check was performed by computing AUC-ROC between original ATAC training input data (dt2) and predicted ATAC. Before ROC computation, matrices where transform into binarized numeric vectors. The threshold employed for binarization is 0.25. This computation was performed using *roc and auc* functions from *pROC* package.

#### *Default model development.*

Computing scores: Neural networks have a certain degree of variability while learning a model; therefore, the results shown have been generated using the mean of 10 trained networks for the case of LIBRA, Autoencoder, Variational Autoencoder or Beta Variational Autoencoder. The highest performance scores in each set of 10 trained models have been highlighted in appropriate cases.

Hyperparameter selection: In order to generate the best configuration for LIBRA according to the nature of the single-cell data, a series of analyses have been conducted using DataSet1. Among them, hyperparameters, layers, number of nodes, activation function used, or optimizer applied has been tested over a basic structure presented in Table S1; as a result, the final framework is shown in Fig.S1A,B.

(i) *Autoencoder-type configuration* ("Selection of Autoencoder Model" table): several neural network models have been investigated. Results show that models based on distribution fits such as VAE or BVAE provide a lower model quality (higher values of Q1 and Q2) but also return worse granularity on cell sub-type identification (lower values of PPJI). As a result, we selected the AE architecture for the NN1 model.

(ii) *The number of dimensions of the projected space* ("Selection of # of middle layer neurons" table): we observed that for a paired sample of about 5,000 to 10,000 cells the highest performance in biological preservation (PPJI) was obtained when considered 5 to 10 dimensions. Such selection has as an additional trade-off: a worse model quality (high Q1) but better dt2 encoding (lower Q2). We selected ten as the number of dimensions in the middle layer.

(iii) *Selection between peak and gene-activity derived information for ATAC-seq*: we observed that the use of the original peaks matrix information from scATAC outperforms the use of the transformed activity expression when considering PPJI (cell sub-type identification) and model quality (Q1). The trade-off is a worse dt2 encoding (Q2).

(iv) *The assignation of dt1 and dt2* ("Selection of the order" table): we observed that setting dt1=ATAC and dt2=RNA returns better PPJI (cell sub-type identification) and model quality (Q1); as a trade-off dt2 encoding (Q2) is worse. We consider that an explanation for this is the different levels of granularity derived from scRNA and scATAC.

(v) *The features to be included* ("Selection of features to include: all vs most variable" table): the use of only the MVF shows a better model quality (Q1) and dt2 encoding (Q2). When comparing cell sub-type identification (PPJI), we observed that on average "all features" has better results, but the "MVP" returns the best model from 10 runs. Signac[4] library has been used for selecting among different percentages of most variable peaks obtaining a worse performance (data not shown).

(vi) *The number of intermediate layers*: we compared using one or two hidden layers. When using 2 hidden layers, we observed improvement for all quality measures Q1, Q2 and PPJI.

#### *LIBRA structure and additional hyperparameters.*

LIBRA neural networks were implemented by using Keras R and Python packages. The final structure used for LIBRA consists of two neural networks; the primary/first one (NN1) implemented to identify the shared latent space (SLS). The second one (NN2) is used for generating a mapping (encoding) of dt2 into the SLS modalities. NN1 contains two parts the encoder and decoder ones while NN2 only includes an encoder. The characteristics of the networks and encoding are:

- The number of layers and nodes used for each of these networks are shown in Fig.S1(a,b).
- Because of the sparse nature of single-cell data, Leaky ReLU has been used as an activation function to avoid zero components in the middle layer, making middle layer size an attribute that can be modified based on the number of paired cells used on the data.
- Adam has been used as an optimizer, and the loss function used is the mean squared error (MSE). Adam optimizer has been configured with; 0.001 learning rate, 0.9 for beta_1 and 0.999 for beta_2 values. The values for epsilon was set to the default, i.e., 0.001 and decay to 0.
- A dropout rate of 0.2 has been used for all activation functions except for the last to the output layer where no dropout is used.
- The maximum number of epochs was set at 1.500, but an early stopping rule with a patience value of 20 over loss score was used, causing the learning process to stop if no improvement was obtained after 20 epochs. We used a large batch size of 7000 to speed up computations on CPU-based systems (limited parallelization capabilities), while smaller datasets will use the maximum available samples; otherwise, BABEL has been fixed to 512 batch size, because it was trained on a GPU (NVIDIA Tesla V100 with an excellent parallelization capabilities) reducing epoch training time and the number of epochs required to achieve the convergence.

- Learning rate plateau callback was added with a factor value of 0.1, starting from 0.001 and updating it with a patience of 15 until reaching a learning rate of 0.00001.

When using LIBRA, the following applies:

- The components generated in the middle layer of the main LIBRAS's neural network (NN1) were used for the Louvain clustering application and, therefore, cluster identification. The resolution used is 0.8 and k=30, in the same way as before the clustering was repeated over 1,000 different combinations of starting points to have a robust clustering outcome.
- UMAP dimensionality reduction method was applied for visualization generation over the middle layer obtained components.

(vii)Q1, Q2 and PPJI.

##### *LIBRA fine-tune (deep adaptive structure).*

We change the static default model to a dynamic one based on the data. To do this, we perform an exhaustive search using a grid of hyperparameters ranging as: number of layers [1,2,3,4,5,6], number of nodes [256,512,1024,2048], alpha [0.1,0.3,0.5], dropout [0.1,0.2,0.3,0.4] and mid layers size [10,50,70] for integration task and number of layers [1,2], number of nodes [128,256,512], alpha [0.05,0.1,0.3], dropout [0.1,0.2], batch size [32,64,128] and mid layers size [10,30,50,70].

To increase the computation speed, it has been implemented in parallel to train multiple models at the same time.

A convex decreasing curve was used for number of nodes assignation to layers to avoid reduce it linearly based on number of layers used. This was done dividing the maximum number of nodes used by a factor of “2 * layer position” for a given layer; Example: for a 2048 number of nodes hyperparameter having 6 layers nodes of encoder will by assigned as, 2048 (1° intermediate layer), 1024 (2° intermediate layer), 512 (3° intermediate layer), 341 (4° intermediate layer), 256 (5° intermediate layer) and 204 (6° intermediate layer). By doing this we allow a high increase of the layers without the need to assign an extremely high value in the first layers so as not to suffer from a negligible value of nodes in the last ones, which would happen if we used a linear decrement Example: dividing by 2 based on previous layer size.

In our tests python implementation of LIBRA has shown to be able to train the model significantly faster than in R outcomes obtained, python version requires less resources (RAM). For this reason, it has been implemented in both to extend access to users and take advantage of python properties.

##### *Parameters for the integrative tools:*

BABEL[12]: We used the implementation from https://github.com/wukevin/babel with default parameters for all datasets. Same normalized data input used for the training of the rest of the models was used as input, generating a BABEL shared space of 16 components. Adam optimizer was used, with an initial learning rate of 0.01 and a batch size of 512 were set by default. Shared space generated by BABEL was used for clustering computation and PPJI estimation. NVIDIA Tesla V100 GPU with 32 GB RAM was used for the training and inference of BABEL on all datasets.

totalVI[11]: The publicly available implementation from totalVI at https://docs.scvi-tools.org/en/stable/user_guide/notebooks/totalVI.html has been used. Default model values have been selected generating a shared space of a total of 10 components. As guidelines present, count data from already filtered datasets (same input for all models) has been used as an input for totalVI by using also the 2.000 mvg for transcriptomic data. In order to compare the integration quality with the rest of the models, shared space generated has been used for clustering and PPJI computation.

MOFA+[10]: As for the rest of the models tested, corresponding guidelines present at https://github.com/bioFAM/MOFA have been used. A maximum of 15 factors has been generated in the shared space using MOFA+ for the different datasets employed. MOFA+ reduced factors' space has later been used for integration performance measurement by using PPJI metric after clustering computation over this space.

##### *CITE-seq additional quality measurement*

To estimate the level of integration quality for DataSet2, CITE-seq-GSE128639, an additional quality metric has been investigated.

First the nearest cells to each cell have been obtained by applying the k-nearest neighbours algorithm (k-NN) with k=20 over the integrated shared space components from the corresponding model applied (LIBRA, Seurat4, MOFA+, totalVI and BABEL). Then, the mean of the expression values of the 20 cells closest to each cell for each of the 25 proteins was calculated and next the Spearman correlation and Pearson correlation were computed between these generated values for each of the 25 proteins and the original protein expression values.

#### GitHub Repository

The code to reproduce the generation of LIBRA models and the entire analysis described in the manuscript is available at: https://github.com/TranslationalBioinformaticsUnit/LIBRA. It includes a Jupyter notebook and a RMarkDown file.

#### Supplementary Figure Legends.

**Figure S1. LIBRA optimization using DS1.** A. Final configuration of NN1 in LIBRA. B. Final configuration of NN2 in LIBRA. C. Example of bimodal distribution identified when computing Q2 values per cells. Q2 is associated with NN2 quality. D. Pairwise Jaccard Index between the clusterings derived from Seurat4 and LIBRA. E. Heatmap where the color denotes the PPJI distance between the clusters generated using the integrated space of each methodology.

**Figure S2. LIBRA and the identification of additional markers.** A. For each cluster identified in Seurat4 (rows), each line denotes a gene and purple denotes if LIBRA also identifies the gene. B. For each cluster identified in Seurat4 (rows), each line denotes a gene and, if it is identified in the Seurat4 cluster, by LIBRA or by both.

**Figure S3. Integrative analysis in DS4 for Seurar4, BABEL and LIBRA.** Two upper panels to denote the UMAP projection and their clustering for RNA and ATAC, respectively. Then, the following rows are associated to the integrative analysis for the different methods: LIBRA, BABEL and Seurat4. In each row, the right panel shows the integration based clustering, and the two left panels show the integrated based clustering "projected" into the RNA and ATAC UMAPS, respectively.

#### Supplementary Table Legends.

**Table S1. Fine-tuning of LIBRA using DS1: all measures.** Several sub-tables showing Q1, Q2 and PPJI used for the selection of the hyperparameters of LIBRA. In green, the selected version in each case is highlighted. Selections made in upper tables are applied to lower tables. In a large majority of the cases we identified an increased number of clusters in the integrated space.

**Table S2. Fine-tuning of LIBRA using DS1: score.** Scores for each training for each of the fine-steps. Same ordering as table S1, but introducing the weighted *score* for each training. In green the top values are provided.

**Table S3. Robustness and Sensitivity analysis in DS1.** Tables showing Q1, Q2 and PPJI used for the sensitivity and robustness analysis of LIBRA. (a) Quality analysis when pairing information is not correct for different percentages of cells. (b) Quality analysis when dropout is added. (c) Quality analysis for different number of epochs in the training of the model. In a large majority of the cases we identified an increased number of clusters in the integrated space.

**Table S4**. **Number of cells per data-set.**

**Table S5. PPJI values between Seurat 4 and LIBRA comparing the two strategies for normalization.** SCT stands for the sctTransform normalization procedure implemented in Seurat package.

**Table S6**. **Prediction-based comparison between BABEL and LIBRA.** (*) Not conducted in BABEL. (**) DS5 was analyzed using all genes instead of MVG. BABEL framework exceeded the limiting time of 1 week for running on a GPU infrastructure. (***) DS6 was analyzed using up to 2TB ram but greater resources are required for BABEL.

**Table S7. predRNA values for all cells and per cluster in DataSet1**. Rho and p-value denotes the Spearman correlation and significance between the number of cells and *predRNA* computed.

**Table S8**. **predRNA values for all cells and per cluster in DataSet5**. Rho and p-value denote the Spearman correlation and significance between the number of cells and *predRNA* computed.

#### REFERENCES


1. Chen S, Lake BB, Zhang K. High-throughput sequencing of the transcriptome and chromatin accessibility in the same cell. *Nat Biotechnol*. 2019;37(12):1452-1457. doi:10.1038/s41587-019-0290-0

2. Stoeckius M, Hafemeister C, Stephenson W, et al. Simultaneous epitope and transcriptome measurement in single cells. *Nat Methods*. 2017;14(9):865-868. doi:10.1038/nmeth.4380

3. Hao Y, Hao S, Andersen-Nissen E, et al. Integrated analysis of multimodal single-cell data. *bioRxiv*. Published online January 1, 2020:2020.10.12.335331. doi:10.1101/2020.10.12.335331

4. Zhu C, Yu M, Huang H, et al. An ultra high-throughput method for single-cell joint analysis of open chromatin and transcriptome. *Nat Struct Mol Biol*. 2019;26(11):1063-1070. doi:10.1038/s41594-019-0323-x

5. Ma S, Zhang B, LaFave LM, et al. Chromatin Potential Identified by Shared Single-Cell Profiling of RNA and Chromatin. *Cell*. 2020;183(4):1103-1116.e20. doi:https://doi.org/10.1016/j.cell.2020.09.056

6. Clark SJ, Argelaguet R, Kapourani C-A, et al. scNMT-seq enables joint profiling of chromatin accessibility DNA methylation and transcription in single cells. *Nat Commun*. 2018;9(1):781. doi:10.1038/s41467-018-03149-4

7. Stuart T, Butler A, Hoffman P, et al. Comprehensive Integration of Single-Cell Data. *Cell*. 2019;177(7):1888-1902.e21. doi:10.1016/j.cell.2019.05.031

8. Stuart T, Srivastava A, Lareau C, Satija R. Multimodal single-cell chromatin analysis with Signac. *bioRxiv*. Published online January 1, 2020:2020.11.09.373613. doi:10.1101/2020.11.09.373613

9. Waltman L, van Eck NJ. A smart local moving algorithm for large-scale modularity-based community detection. *Eur Phys J B*. 2013;86(11):471. doi:10.1140/epjb/e2013-40829-0

10. Argelaguet R, Arnol D, Bredikhin D, et al. MOFA+: a statistical framework for comprehensive integration of multi-modal single-cell data. *Genome Biol*. 2020;21(1):111. doi:10.1186/s13059-020-02015-1

11. Gayoso A, Steier Z, Lopez R, et al. Joint probabilistic modeling of single-cell multi-omic data with totalVI. *Nat Methods*. 2021;18(3):272-282. doi:10.1038/s41592-020-01050-x

12. Wu KE, Yost KE, Chang HY, Zou J. BABEL enables cross-modality translation between multiomic profiles at single-cell resolution. *Proc Natl Acad Sci*. 2021;118(15):e2023070118. doi:10.1073/pnas.2023070118

13. Tang M, Kaymaz Y, Logeman BL, et al. Evaluating single-cell cluster stability using the Jaccard similarity index. *Bioinformatics*. Published online November 9,

2020. doi:10.1093/bioinformatics/btaa956
